# Teleost-specific ictacalcins exhibit similar structural organization, cation-dependent activation and transcriptional regulation as human S100 proteins

**DOI:** 10.1101/2025.06.10.658860

**Authors:** Liz Hernández, Théo Paris, Maria Demou, Catherine Birck, Christina Begon-Pescia, Juan Francisco Rodríguez-Vidal, Sylwia D. Tyrkalska, Charlotte Bureau, Catherine Gonzalez, Juliette Gracia, Etienne Lelièvre, Victoriano Mulero, Mai Nguyen-Chi, Laure Yatime

## Abstract

S100 proteins are highly versatile calcium-binding proteins from vertebrates. Following extracellular release, they become instrumental in immune and antimicrobial defenses, initiating the inflammatory response through receptor signaling and providing direct control of bacterial invaders via nutritional immunity. While mammalian S100s have been extensively studied, very little is known about the more recently discovered S100 proteins from teleost fish, including those with no strict orthologs in mammals. Comparable functioning between both clades would allow to expand their study into the highly popular zebrafish model, particularly suited for live imaging and mechanistic exploration of immune and inflammatory processes. To fill the gap of knowledge on teleost S100s, we here provide a detailed structural and biochemical characterization of S100i1 and S100i2 from *Danio rerio*, two teleost-specific S100s absent in mammals. We demonstrate that they nevertheless share conserved tertiary and quaternary organization with mammalian S100s. In addition, they exhibit comparable calcium binding properties and undergo a similar calcium-dependent activation mechanism. Furthermore, they display analogous expression pattern, being enriched in tissues highly exposed to the environment like gills and skin, the latter constituting an important reservoir of S100 proteins in mammals. Finally, our results show, for the very first time, that *s100i2*/*i2* gene expression is differentially modulated in sterile disease conditions associated with sustained inflammation or high hypoxic state. Altogether, these findings underline the strong parallelism existing between mammalian and teleost-specific S100 proteins despite their divergent evolution, opening up new avenues to explore their biology in the zebrafish model.

## Introduction

S100 proteins are small peptides of 10-15 kDa that form the largest sub-group within the EF-hand superfamily of calcium-binding proteins. They are exclusively found in vertebrates and exert pleiotropic functions, both inside and outside cells, acting as calcium-dependent regulators of many vital processes, such as metabolism, cell proliferation and cytoskeleton rearrangement within cells, or immune and antibacterial defenses in the extracellular milieu [1,2]. Noteworthily, several S100 members are considered as major effectors of innate immunity in humans. In response to infections or sterile injuries, they are released in the extracellular environment, where they contribute both to the propagation of inflammation, through their alarmin function, and to the antimicrobial response, through nutritional immunity [3,4]. Their role in inflammation is, however, double-edged, as long-lasting injuries allow for strong up-regulation of their gene expression, thus promoting their continuous release. This feeds the inflammatory response up to a non-resolving state that leads to aggravated damage and, ultimately, to chronic pathologies [5]. Hence, master regulators of inflammation like the S100A8/A9 heterodimer are considered both as valuable biomarkers and promising therapeutic targets in numerous disease conditions linked to aberrant inflammation [6].

Given the importance of S100 proteins in human physiopathology, animal models to study their function in health and disease have long been developed, predominantly in rodents. Until the last decade, S100 proteins were thought to be restricted to mammals, in which around twenty members are commonly annotated. More recent phylogenetic analyses revealed the presence of orthologous genes in distinct vertebrate clades, including amphibians and teleost fish [7]. Among the latter, zebrafish has become a highly popular animal model to study immune and inflammatory processes, having a high degree of genetic and immune conservation with mammals and offering selective advantages as compared to mice models, notably its transparency at larval stages that allows to follow physiological processes in real time thanks to intravital imaging [8,9]. Despite such attractive traits, zebrafish has so far not been considered extensively for the study of S100 protein-related processes, and in fact, still very little is known about the expression and function of zebrafish S100 family members [10].

Interestingly, a distinct set of *s100* genes is found in teleost fish, that share no evident phylogenetic lineage with any mammalian *S100* genes, which questions their conservation in terms of structural and functional features [7,11]. The first teleost-specific gene was discovered in the chemosensory tissues of catfish (*Ictalurus punctatus*) and was therefore termed ictacalcin (*icn*) [12]. Two paralogs of this gene, that arose from the third whole genome duplication event, were more recently identified in zebrafish: *icn* and *icn2*, also referred to as *s100i1* and *s100i2* for consistency with the S100 family nomenclature [13]. While these two fish *s100* genes have been studied in slightly more details due to their predominant expression in epithelial tissues, the degree of resemblance of their products with their mammalian counterparts, and thus the possible parallelism in biological functions, remains to be clarified.

To gain insights on how zebrafish-specific S100 proteins compare to mammalian ones, we here provide a detailed structural and biochemical characterization of the two ictacalcin proteins from *Danio rerio*, combining X-ray crystallography and isothermal titration calorimetry. Using the zebrafish model, we further explore ictacalcin gene expression under homeostatic conditions and in various sterile disease models, revealing *s100i1*/*i2* transcriptional modulation in conditions associated with sustained inflammation or high hypoxic state. Altogether, these findings highlight the strong similarities existing between mammalian and teleost-specific S100 proteins, hinting for possible analogous functions despite their divergent evolution.

## Results

### Zebrafish ictacalcins possess conserved tertiary and quaternary organization with mammalian S100 proteins

To assess the structural properties of zebrafish ictacalcins, we produced them as recombinant proteins and crystallized them in various ionic conditions, yielding three distinct crystal forms that all diffracted to fairly high resolution (Table 1) and which we attributed to apo S100i1 (crystal form 1), Na^+^-bound S100i1 (crystal form 2) and Mg^2+^-bound S100i2 (see later). The final atomic models, covering residues Ala2 to Thr92/Gly93 for S100i1 and residues Met5 to Phe92 for S100i2, are displayed in Figure 1, together with final electron density maps. At the protomer level, both S100i1 and S100i2 adopt the classical four-helix bundle fold characteristic of S100 proteins (Figs. 1A, 1C & 1E). The four helices are formed by the same stretches of residues in both proteins, corresponding in S100i1 to Ser4-Ser21 (helix H1), Ser31-Leu43 (helix H2), Asp52-Asp64 (helix H3) and Asp72-Leu86 (helix H4). A short α-helix turn (H2’) is also observed both in S100i1 and S100i2, in the hinge region linking helices H2 and H3, that covers residues Gly44 to Phe47 (S100i1 numbering), and another short α-helix turn (H4’) is present after helix H4 in the apo S100i1 structure, encompassing residues Glu89 to Thr92. The last five C-terminal residues were not visible in the density, most probably corresponding to flexible regions without defined secondary structure.

**Figure 1.**
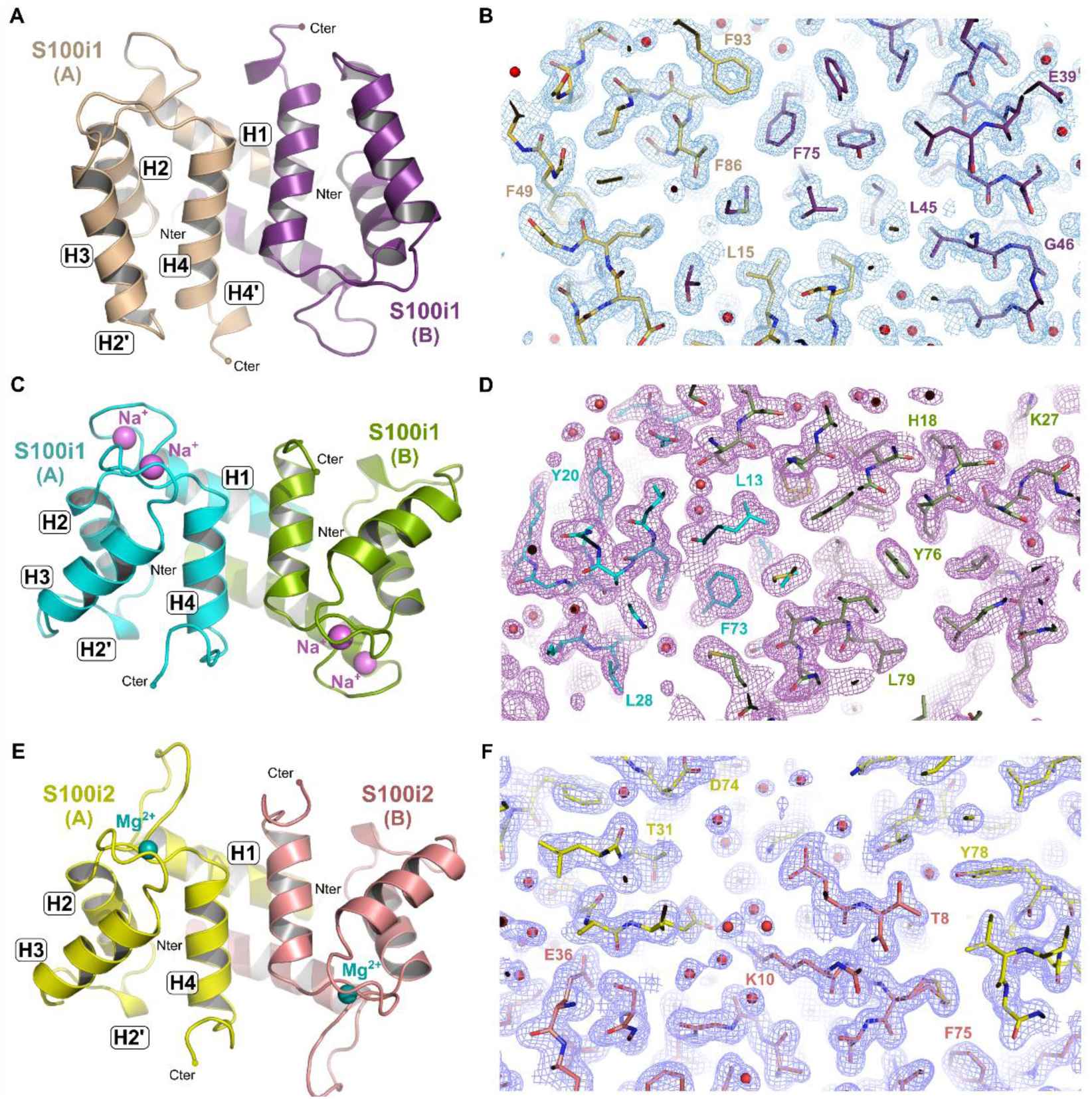
Crystallographic structures of zebrafish ictacalcins in various ionic states. **A.** General overview of the structure of apo S100i1 homodimer at 1.5 Å resolution. The two protomers are shown in beige and purple. **B.** Final electron density map contoured at 1σ (blue mesh) with final model superimposed. **C.** General overview of the structure of Na^+^-bound S100i1 homodimer at 1.65 Å resolution. The two protomers are shown in cyan and olive. Sodium ions are displayed as magenta spheres. **D.** Final electron density map contoured at 1σ (purple mesh) with final model superimposed. **E.** General overview of the structure of Mg^2+^-bound S100i2 homodimer at 1.6 Å resolution. The two protomers are shown in yellow and salmon. Magnesium ions are displayed as blue spheres. **F.** Final electron density map contoured at 1σ (purple blue mesh) with final model superimposed. For each structure, helix nomenclature is indicated on protomer A.

**Table 1.**
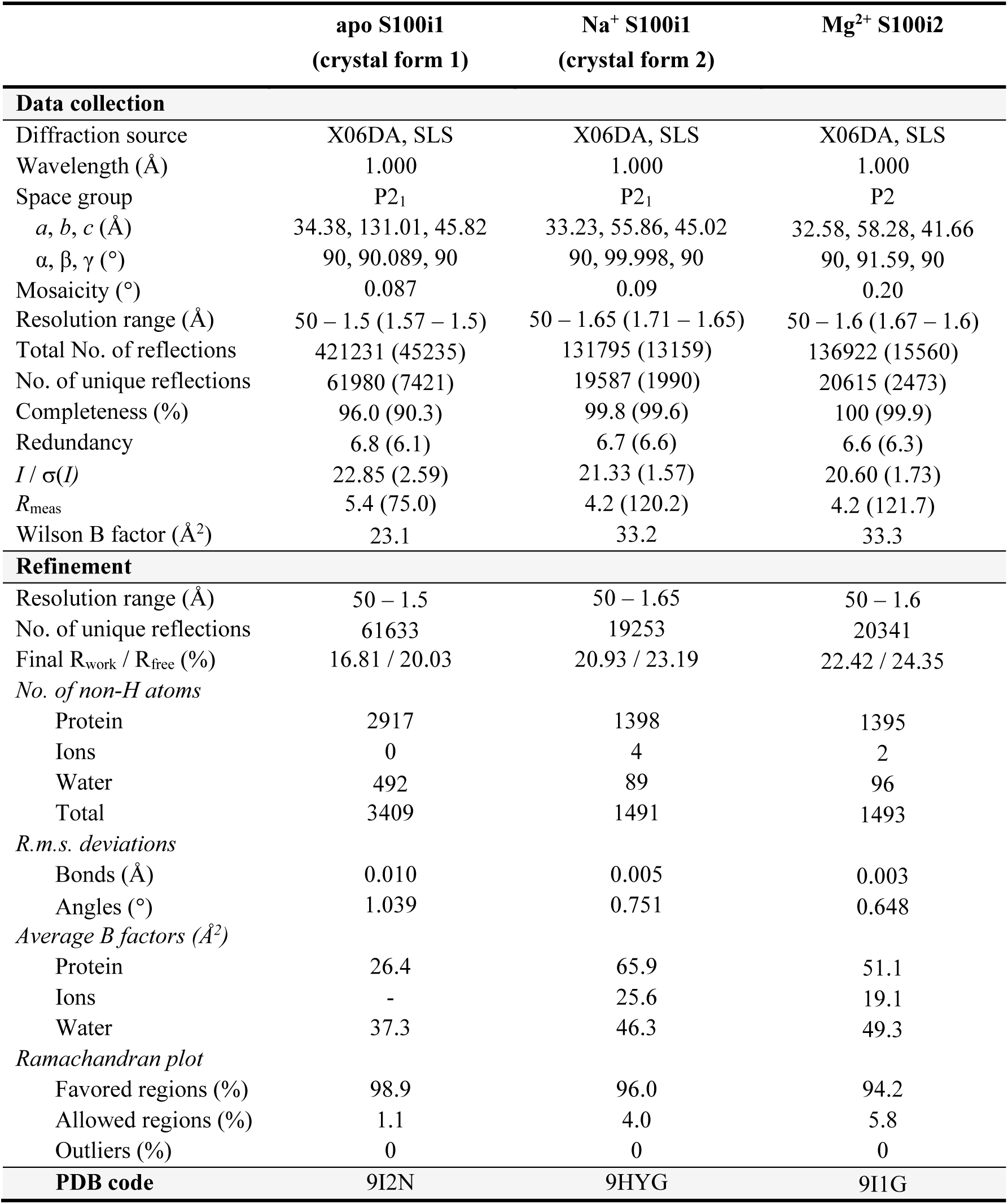
Data collection and refinement statistics. Values indicated in parentheses correspond to the last shell of resolution.

As expected for S100 proteins, and in agreement with their elution profile on size exclusion chromatography (Figs. 2A & 2E), both ictacalcins organize into homodimeric assemblies that follow the canonical H1-H1-H4-H4 packing seen in most S100 structures (Figs. 2B & 2F). In both proteins, the dimer interface is held in place by an extended network of hydrophobic interactions involving the side chains of aromatic and aliphatic residues carried by the central portions of H1 and H4 from both protomers (Figs. 2C & 2G). The interface is further stabilized at both extremities by a smaller network of hydrogen bonds involving side chains of polar residues at the N-terminus of H1 from one protomer, and at the hinge region (H2-H3 linker) and C-terminus of H4 from the second protomer, either through direct contacts such as Ser4-Glu42, Thr6-Glu42 or Gln7-Asp45 (S100i1 numbering), or through water-mediated inter-protomer interactions (Figs. 2D & 2H). The interface areas calculated by PISA [14] cover 1105 Å^2^ for S100i1 and 814 Å^2^ for S100i2, with 34 and 37 interfacing residues, respectively, and 11 hydrogen bonds in both cases. The overall solvation free energy gain is of -21.4 and -15.4 kcal/mol for s100i1 and s100i2 dimers, respectively, with associated P-values of 0.114 and 0.043, which highlights the strong hydrophobicity and stability of the interfaces. Altogether, this demonstrates that zebrafish ictacalcins adopt a quaternary organization identical to that of mammalian S100s.

**Figure 2.**
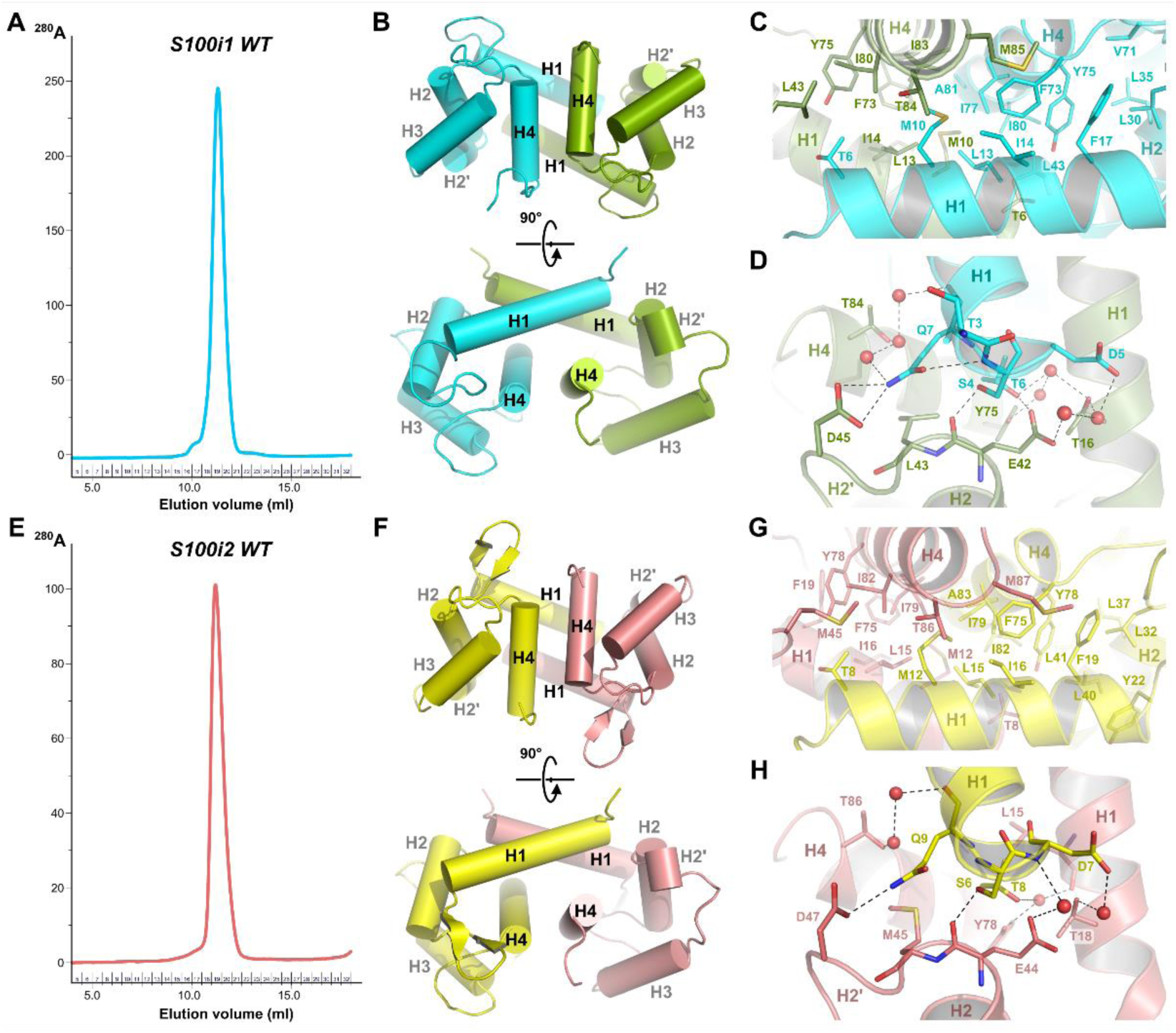
Homodimeric organization of zebrafish ictacalcins. **A.** SEC elution profile of WT S100i1 on Superdex 75 Increase, revealing predominant elution as a homodimer. **B.** Helix packing in the S100i1 homodimer. **C.** Hydrophobic interactions stabilizing the S100i1 dimer interface. **D.** Electrostatic interactions stabilizing the S100i1 dimer interface. **E.** SEC elution profile of WT S100i2 on Superdex 75 Increase, revealing predominant elution as a homodimer. **F.** Helix packing in the S100i2 homodimer. **G.** Hydrophobic interactions stabilizing the S100i2 dimer interface. **H.** Electrostatic interactions stabilizing the S100i2 dimer interface.

### Ictacalcins display conserved cation binding mode in their EF-hand motifs

The presence of various cations in the crystallization buffers allowed to trap S100i1 and S100i2 in several ionic states. Hence, while the EF-loops of S100i1 in crystal form 1 are devoid of cations (Fig. 3A), two Na^+^ ions are found in each s100i1 protomer of crystal form 2, occupying both EF-loops of the protein (Fig. 3B). Additionally, one Mg^2+^ ion is found in the second EF-loop (EF2; H3-H4 linker) of each S100i2 protomer, whereas the first EF-loop (EF1; H1-H2 linker) remains empty (Fig. 3C). Comparison of the EF-loop conformations in both apo and ion-bound states reveals important conformational rearrangements, as for mammalian S100s. The two EF-loops shift from a rather extended conformation when empty, to a more compact architecture when occupied, wrapping up around the central ion as classically observed in Ca^2+^-loaded human S100 proteins (Fig. 3D).

**Figure 3.**
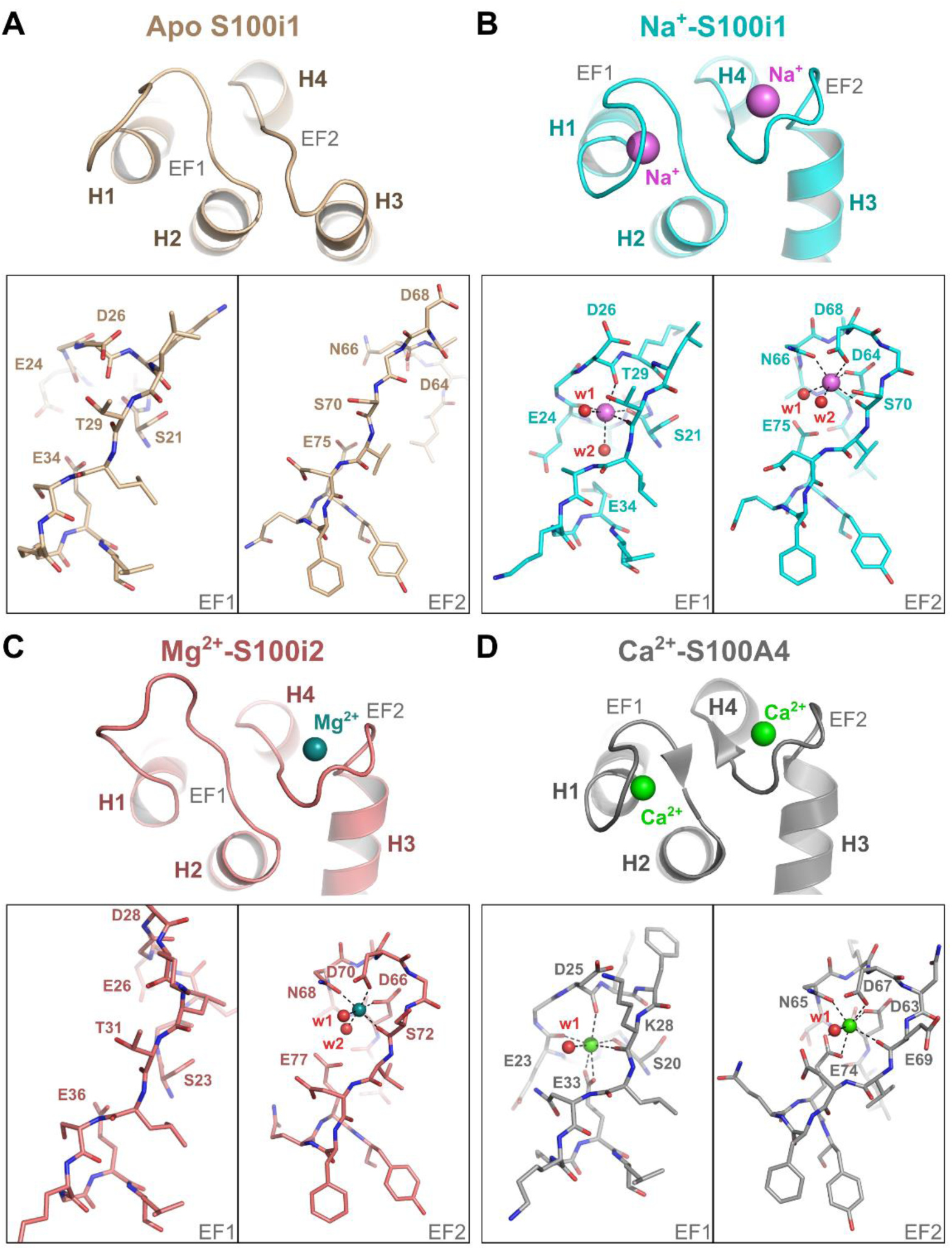
Cation binding in the EF-loops of S100i1 and S100i2. **A-C.** Close-up view on the overall conformation and cation binding residues within the EF-loops of apo S100i1 (**A**), Na^+^-bound S100i1 (**B**) and Mg^2+^-bound S100i2 (**C**). **D.** Comparison with the EF-loops of Ca^2+^-bound human S100A4 (PDB_ID 2Q91 [52]).

These structures also reveal the residues involved in cation binding in both EF-hand motifs, the average ion-oxygen distances being of 2.43 Å for Na-O bonds and 2.06 Å for Mg-O bonds, in agreement with mean values reported in the literature [15,16]. Sodium displays an octahedral coordination sphere in both EF1 and EF2, with hydrogen bonds provided by main chain carbonyls of Ser21, Glu24, Asp26, Thr29 (EF1) and Ser70 (EF2), side chain oxygens of Asp64, Asn66 and Asp68 (EF2) and two water molecules in each EF-loop (held in place by Glu75 in EF2). Magnesium has similar octahedral coordination geometry in EF2 of S100i2, with the same residues implicated in cation chelation, namely Asp66, Asn68 and Asp70 through side chain oxygens, Ser72 through main chain carbonyl and two water molecules, brought by Glu77. Like in mammals, the two S100i1/i2 EF-loops are thus non-equivalent, the second one providing coordination mainly via side chain oxygens, thus corresponding to a canonical EF-loop, whereas the first loop chelates cations via main chain carbonyl, thereby corresponding to a pseudo EF-loop (S100-specific).

Strong conservation of the S100i1/i2 residues involved in Na^+^/Mg^2+^ chelation with those involved in Ca^2+^ coordination in human S100s (Fig. 4A) allows to postulate a conserved calcium binding mode, as illustrated with the 3D-model of Ca^2+^-bound S100i1/i2 (shown in Fig. 4B for S100i1) that we generated. In this model, the EF-loop conformations and overall coordination spheres strongly resemble those observed in Na^+^-S100i1 except that the lateral water molecule at the octahedron basis is now replaced by the two oxygen atoms of Glu34 in EF1 or Glu75 in EF2, thus giving rise to an heptahedral coordination sphere with pentagonal bipyramidal geometry, as observed in Ca^2+^-bound human S100A4 (Fig. 3D).

**Figure 4.**
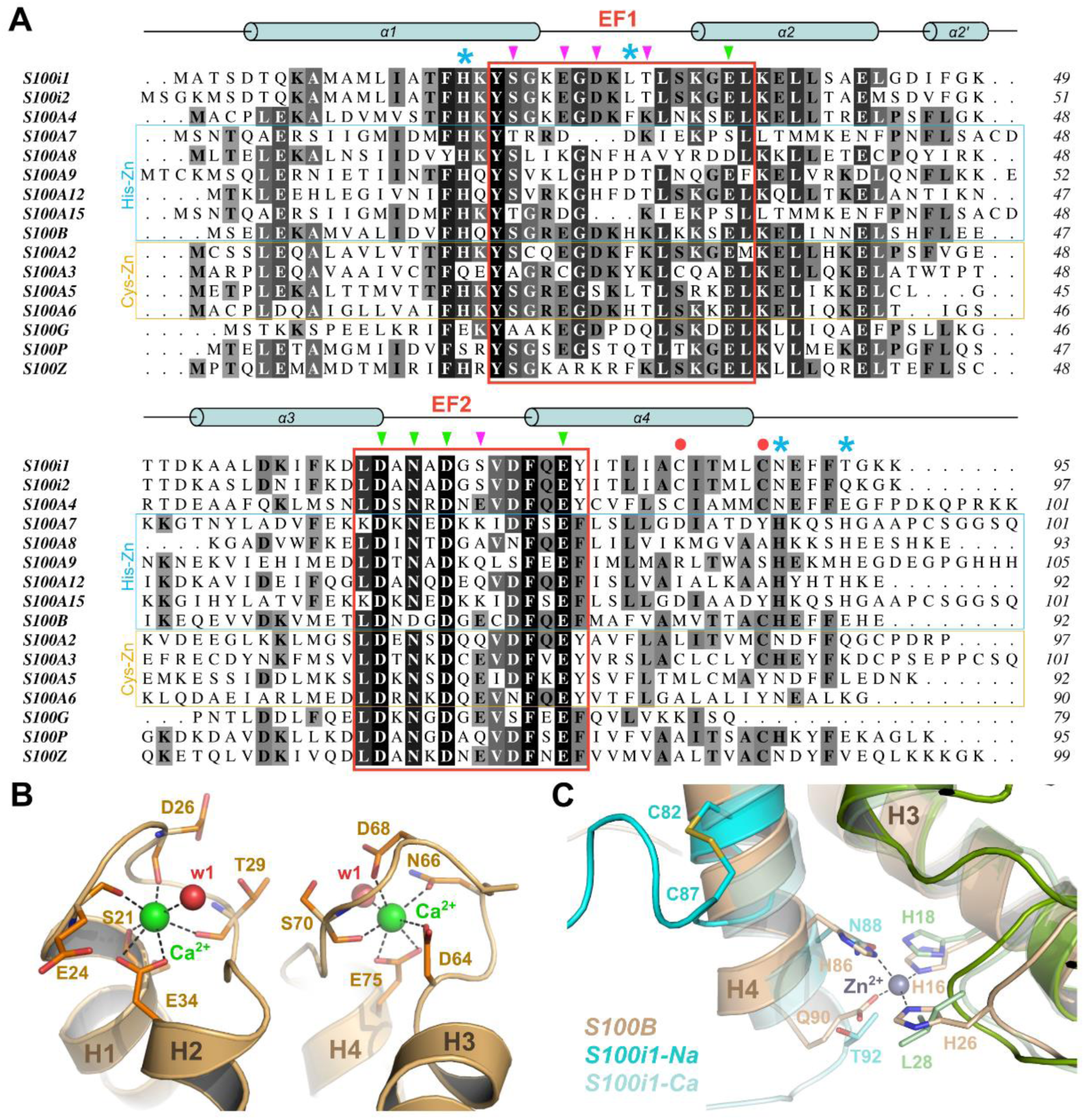
Conservation of the calcium binding mode between zebrafish S100i1/i2 and human S100s. **A.** Sequence alignment of zebrafish S100i1/i2 with human S100s, highlighting the high degree of sequence conservation, especially in the EF-loop motifs (indicated by the red boxes). Residues from EF-loops involved in calcium chelation through main chain or side chain atoms are indicated on top of the alignment with purple or green arrows, respectively. Residues involved in zinc binding in the His-Zn human S100s are indicated with blue asterisks. Cysteine residues involved in intramolecular disulfide bridge in the second half of helix H4 in S100i1/i2 are indicated by a red dot. **B.** AlphaFold-predicted 3D model of Ca^2+^-bound S100i1 highlighting similar binding mode as human S100s. **C.** Superimposition of Zn^2+^-bound human S100B (beige and olive cartoons; PDB_ID 3D0Y [70]) with Na^+^- and predicted Ca^2+^-bound S100i1 showing that the His-Zn site of S100B is not conserved in S100i1. Instead, upon Na^+^ binding at least, the formation of an intramolecular disulfide bridge between Cys82 and Cys87 pushes S100i1 C-terminal tail away from the homodimer interface.

### Cation binding properties of zebrafish ictacalcins

To characterize the binding properties of the calcium sites, we performed isothermal titration calorimetry (ITC) experiments on the two zebrafish ictacalcins. To ensure a better accuracy in the determination of the corresponding dissociation constants (K_D_), we generated CterW mutants encompassing an additional Trp residue at the end of the sequence and thus increasing molar extinction coefficient for a more robust assessment of the protein concentration. Both mutants gave similar production yields, displayed similar SEC elution profiles (i.e. homodimers; Supplementary Fig. S1), and behaved identically as WT proteins. Despite the presence of two binding sites per protomer, a single discernible transition was observed in the ITC titration curves (Figs. 5A-B). Analysis with a macroscopic model of “two symmetric sites” or a microscopic model of “two non-symmetric sites” gave almost similar results but the calculated parameters were better defined using the macroscopic model. The macroscopic K_D1_ and K_D2_ values are respectively 0.99 and 3.96 µM for S100i1 and 1.17 and 4.67 µM for S100i2 (Table 2). For S100i1, the microscopic K_D_ would be 1.98 µM (2 x K_D1_) for both Ca^2+^ binding sites, which is very close to the fitted K_D_ value of 1.99 [0.68 ─ 3.77] µM found for each site using the microscopic model. For S100i2, the microscopic K_D_ would be 2.34 µM for both Ca^2+^ binding sites compared to the fitted K_D_ values of 0.23 [0.003 ─ 5.68] µM and 4.07 [no limit found] µM for each site using the microscopic model. The enthalpies values are much less defined using this latter model and no cooperativity between the two sites was revealed.

**Figure 5.**
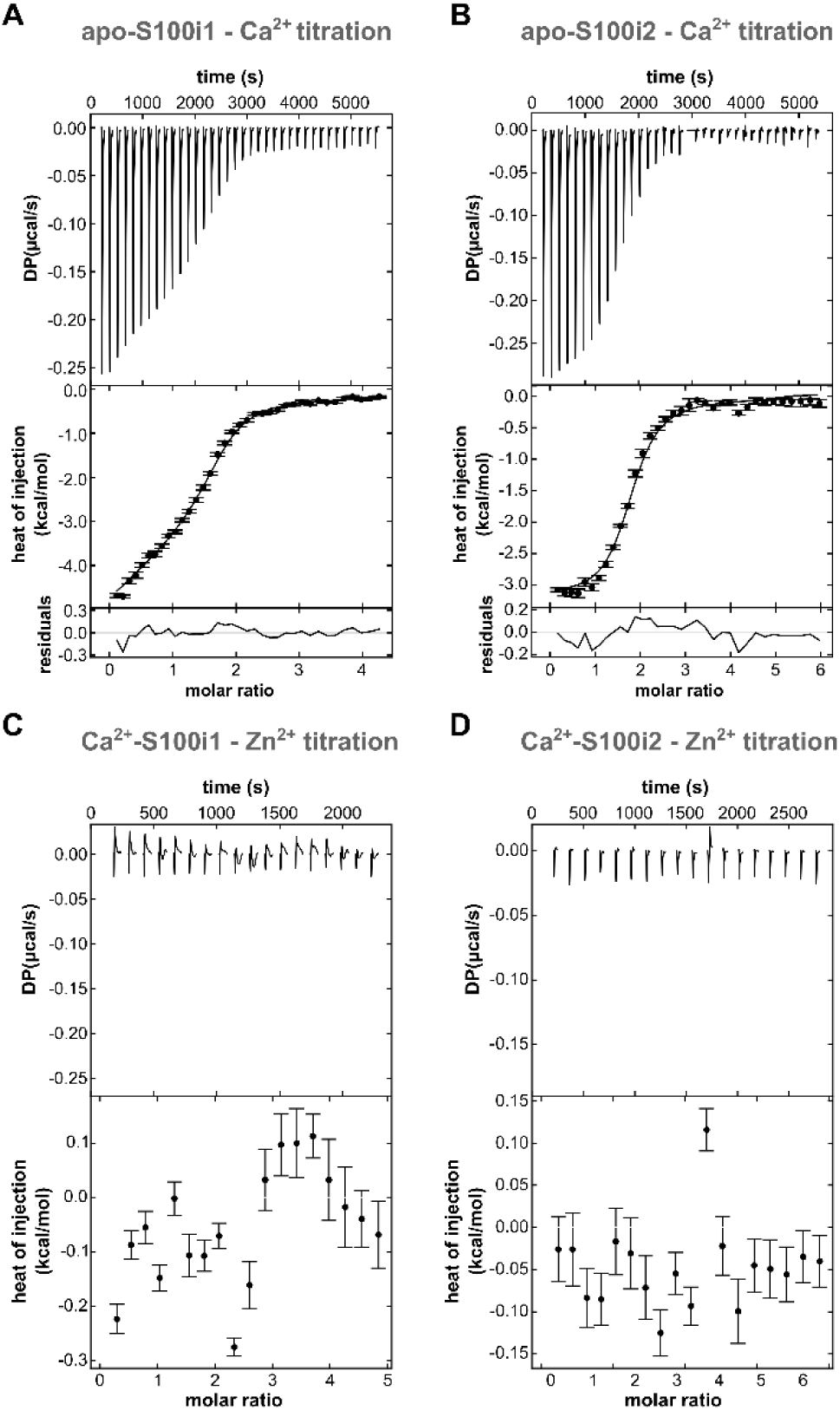
Cation binding properties of zebrafish S100i1/i2. **A-B.** Representative ITC titrations of S100i1 **(A)** or S100i2 **(B)** binding to Ca^2+^. The upper graphs show reconstructed thermograms from NITPIC, the middle graphs show binding isotherms and global fit, and the bottom graphs show the fitting residuals. Shown are titrations of 0.4 mM CaCl_2_ into 40 µM S100i1 and 0.6 mM CaCl_2_ into 42 µM S100i2. In the global analysis, two more titrations were included for S100i1 or S100i2 binding to Ca^2+^, respectively. Macroscopic dissociation constants (K_D_) obtained from global analysis with 2 symmetric sites per protomer are displayed with 95 % confidence intervals in brackets. **C-D.** Representative ITC titrations of S100i1 **(C)** or S100i2 **(D)** binding to Zn^2+^. The upper graphs show reconstructed thermograms from NITPIC and show the corresponding heat changes. Shown are titrations of 1.0 mM Zn acetate into 40 µM S100i1 and 0.6 mM Zn acetate into 60 µM S100i2. No binding to Zn^2+^ can be detected for any of the proteins.

**Table 2.** Dissociation constants and enthalpies for the binding of calcium to zebrafish ictacalcins as measured by ITC. Fitting was performed using a macroscopic model of “two symmetric sites”. Values in brackets indicate 95% confidence interval [2σ] determined by error-surface projection in the global analysis of three experiments

| Protein | K<sub>D1</sub> (μM) | ΔH<sub>1</sub> (kcal/mol) | K<sub>D2</sub> (μM) | ΔH<sub>2</sub> (kcal/mol) | Incompetent fraction (%) |
| --- | --- | --- | --- | --- | --- |
| S100i1 | 0.99 [0.68 – 1.39] | -4.60 [-4.75 – -4.46] | 3.96 | -2.48 [-3.08 – -1.92] | 0.15 [0.10 – 0.19] |
| S100i2 | 1.17 [0.89 – 1.54] | -3.14 [-3.25 – -3.04] | 4.67 | -3.14 [-3.69 – -2.96] | 0.16 [0.13 – 0.19] |

Hence, despite the non-equivalence of the two sites in terms of cation chelating mode, S100i1 and S100i2 bind to calcium with almost equal affinities in the two EF-loops, which is often observed for S100 proteins [17,18].

Several mammalian S100s, notably those involved in antimicrobial defenses through nutritional immunity, also bind transition metals, at sites distinct from the calcium EF-hands [19]. To evaluate the metal binding capacity of S100i1/i2, we next performed zinc titrations in both the apo and Ca^2+^-loaded proteins. No binding could be detected, for neither protein (Figs. 5C-D). This corroborates the fact that only one of the four Zn^2+^-chelating residues present in human S100s is conserved in S100i1/i2, the second His/Asp Zn-ligand in EF1 being instead a leucine, and the two His ligands in the second half of H4 being replaced by an Asn and a Thr/Gln in S100i1/i2 (Fig. 4A). Therefore, as modeled in Fig. 4C in comparison with the Zn^2+^ site of human S100B, the canonical His-Zn site at the interface between the two S100 protomers cannot form in S100i1/i2. Two Cys residues are instead present at the end of H4 in zebrafish ictacalcins but they do not serve as surrogates for metal chelation. Indeed, binding of Na^+^ or Mg^2+^ to S100i1/i2 triggers an unwinding of H4 by one helix turn and leads to the formation of an intramolecular disulfide bridge between these two cysteines (Cys82 and Cys87 in S100i1) that holds the C-terminal tail away from the dimer interface. Whether this S-S bridge also exists in the Ca^2+^-bound form of S100i1/i2 remains to be determined, but it nevertheless seems to prevent favorable positioning of helix H4 C-terminus for the formation of a metal binding site at the dimer interface.

### Ictacalcins undergo cation-dependent conformational rearrangements

Superimposition of our three ictacalcin structures reveals strong conformational movements between the apo form and the two ion-bound models (Fig. 6A). Most notably, H3 undergoes a large rotation of almost 100° upon ion binding, concomitantly with the reorganization of the EF2-loop around the bound ion. The structure of apo S100i1 perfectly superimposes with those of apo S100A2 and apo S100A3 (Fig. 6B), confirming that it represents a genuine apo state of the protein. In contrast, Na^+^-bound S100i1 adopts a conformation very close to that of Ca^2+^- bound S100A2, except for the positioning of H4 that aligns better with H4 of Na^+^-bound S100A12 (Fig. 6C). Importantly, Na^+^-bound S100A12 only contains one Na^+^ ion, in EF2, whereas our Na^+^-S100i1 model bears Na^+^ ions in both EF1 and EF2, which may explain the more distant conformational organization between these two structures. Finally, Mg^2+^-bound S100i2 aligns also quite well with calcium-loaded S100 structures, especially that of Ca^2+^- bound S100A4 (Fig. 6D). Comparing our S100i2 model with that of Mg^2+^-bound S100G, the sole S100 protein structure available with bound magnesium, highlights the intermediary conformation adopted by Mg^2+^-S100i2, with helices H3 and H4 being positioned half-way between those of Ca^2+^-S100A4 and Mg^2+^-S100G.

**Figure 6.**
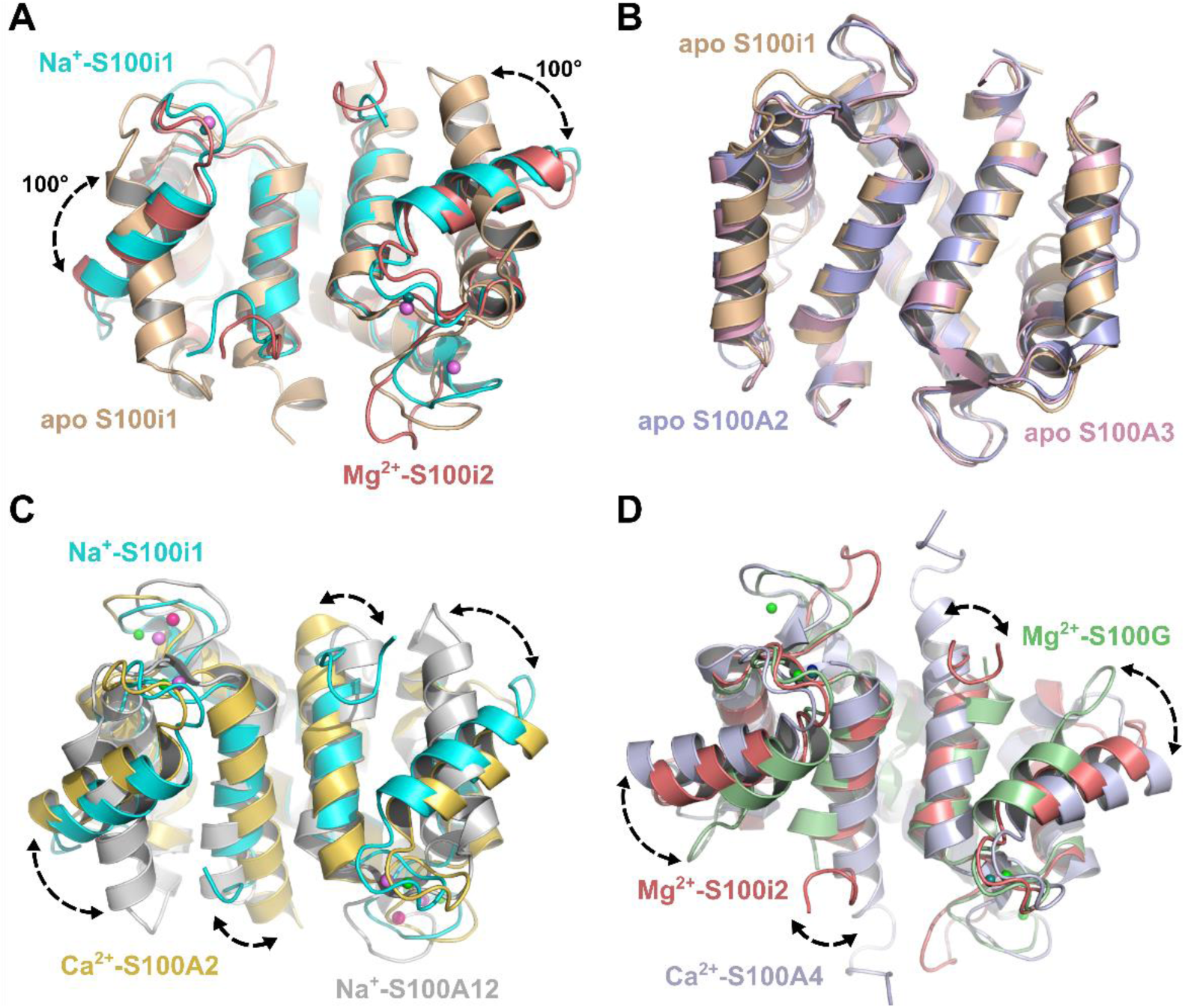
S100i1 and S100i2 undergo conformational rearrangements upon cation binding. **A.** Superimposition of the structures of apo S100i1 (beige), Na^+^-S100i1 (cyan) and Mg^2+^- S100i2 (dark salmon) homodimers highlights major conformational rearrangements upon occupation of the EF-loops by Na^+^ or Mg^2+^. The most prominent one is the reorientation of helix H3 by a rotation of 100°. **B.** Comparison of the overall conformation of apo S100i1 with those of apo human S100A2 (PDB_ID 2RGI [71]) and apo human S100A3 (PDB_ID 1KSO [72]), highlighting that the protein is in a genuine apo conformation in absence of cation bound in the EF-loops. **C.** Comparison of the overall conformation of Na^+^-S100i1 with those of Ca^2+^- bound human S100A2 (PDB_ID 4DUQ [51]) and Na^+^-bound human S100A12 (PDB_ID 2WCE [73]). Na^+^-S100i1 adopts a conformation close to that of Ca^2+^-S100A2, in contrast to Na^+^-S100A12 which resembles more closely the apo conformation of S100 proteins. **D.** Comparison of the overall conformation of Mg^2+^-S100i2 with those of Ca^2+^-bound human S100A4 (PDB_ID 2Q91 [52]) and Mg^2+^-bound human S100G (PDB_ID 1IG5 [74]). Mg^2+^-S100i2 adopts an intermediary conformation between the fully activated Ca^2+^-S100A4 one, and the partially activated Mg^2+^-S100G one.

Thus, binding of Na^+^ or Mg^2+^ to S100i1/i2 induces conformational rearrangements within the protein, predominantly driven by the repositioning of helix H3, and yielding intermediary conformations as compared to Ca^2+^-loaded human S100 structures. These observations suggest that S100i1/i2 do not act as calcium buffers, but instead undergo calcium-dependent conformational activation, like their mammalian counterparts, that may serve as switch on for their interaction with target effectors.

### The spatio-temporal distribution of ictacalcin transcripts strongly parallels that of mammalian *s100* genes

To further evaluate the resemblance between zebrafish S100i1/i2 and mammalian S100s, we analyzed their *in vivo* spatio-temporal distribution, initially under homeostatic conditions. Previous studies already reported partial expression data for *s100i1*/*i2* genes, using either RT-qPCR analyses or *in situ* hybridization [7]. The major bottleneck of these investigations, however, lied in the sets of primers and RNA probes employed, which could not distinguish between *s100i1* and *s100i2*, thus providing global data for *i1*/*i2* expression. To circumvent this, we took great care in designing highly specific sets of primers, choosing them in the regions of most divergence between *s100i1* and *s100i2* coding sequences (Fig. 7A). RT-PCRs performed on plasmids encompassing the fourteen different zebrafish *s100* coding DNA sequences confirmed that our primers are highly specific for the target gene and do not amplify the other duplicate gene nor any of the twelve other zebrafish *s100* genes (Fig. 7B).

**Figure 7.**
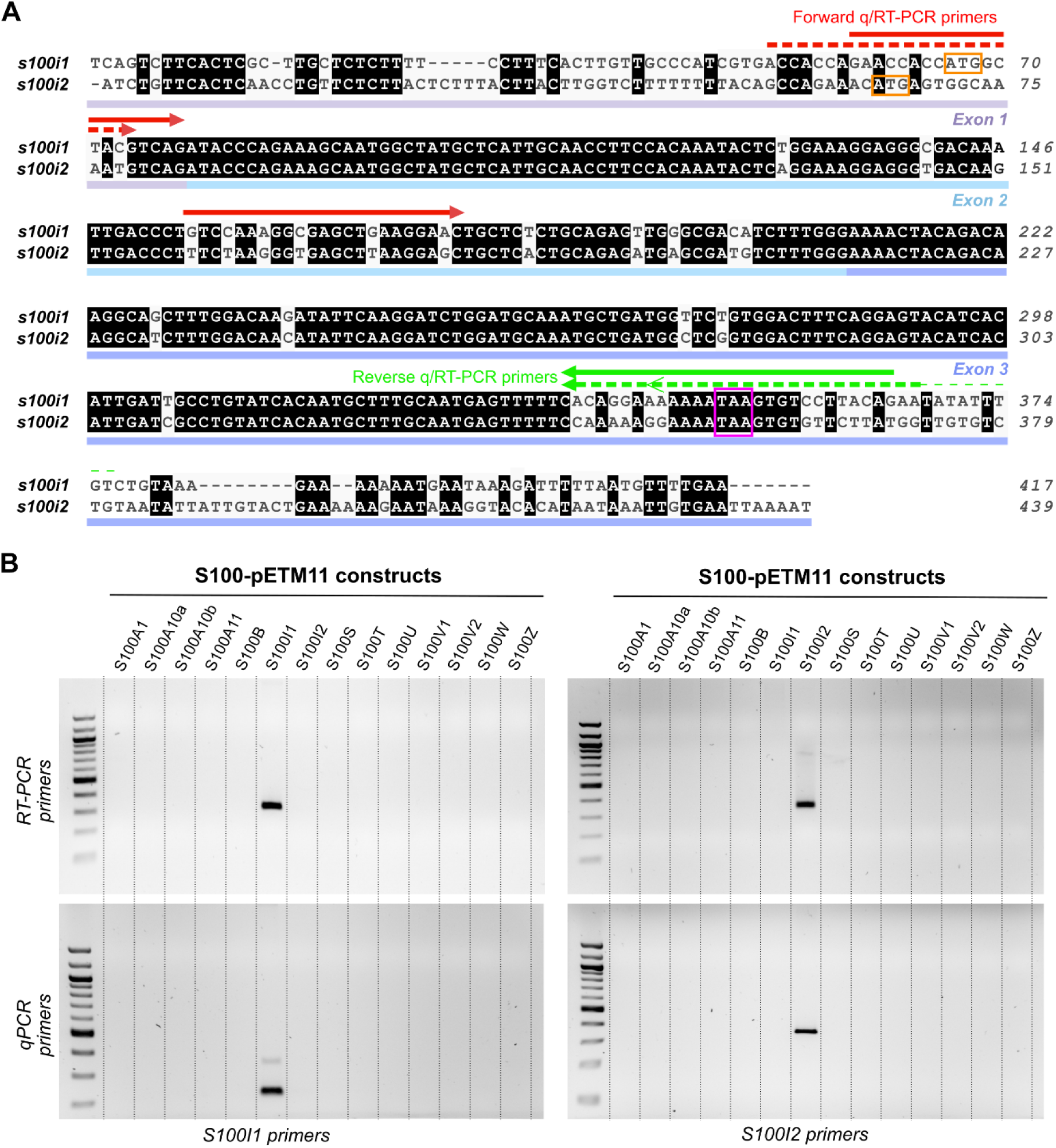
Specificity check for s100i1/i2 RT/qPCR primers. **A.** Alignment of the full-length coding sequences of zebrafish s100i1 and s100i2, colored according to sequence conservation. Positions of the three exons are indicated below the sequences. The start and stop codons are marked by orange and purple boxes, respectively. The forward and reverse primers used for semi-quantitative PCR (RT-PCR) and quantitative (qPCR) are indicated above the alignment (full line for s100i1, dotted line for s100i2; see Supplementary Table S1). **B.** Specificity check on the S100-pETM11 constructs encompassing the fourteen distinct s100 coding sequences. The set of primers chosen for each ictacalcin gene only amplifies the corresponding construct, demonstrating its high specificity. The DNA ladder employed is the 100 bp DNA ladder from New England Biolabs.

Given these highly specific primers, we first investigated when *s100i1*/*i2* start being expressed in zebrafish embryos, by conducting RT-PCRs on total RNA samples collected at different developmental stages. As shown in Figure 8A, both genes are readily detected at 10 hpf and their expression remains stable up to adulthood, suggesting that they are required early during development and then throughout the life. Absence of detection before 10 hpf indicates no or minimal maternal contribution to their expression. Next, we examined *s100i1*/*i2* distribution in eight representative adult tissues/organs. Interestingly, the two duplicate genes display rather distinct expression patterns, *s100i1* being broadly expressed in all samples investigated whereas *s100i2* is mainly restricted to skin and gills, and to a lesser extent to heart, gut, testis and ovaries (Fig. 8B).

**Figure 8.**
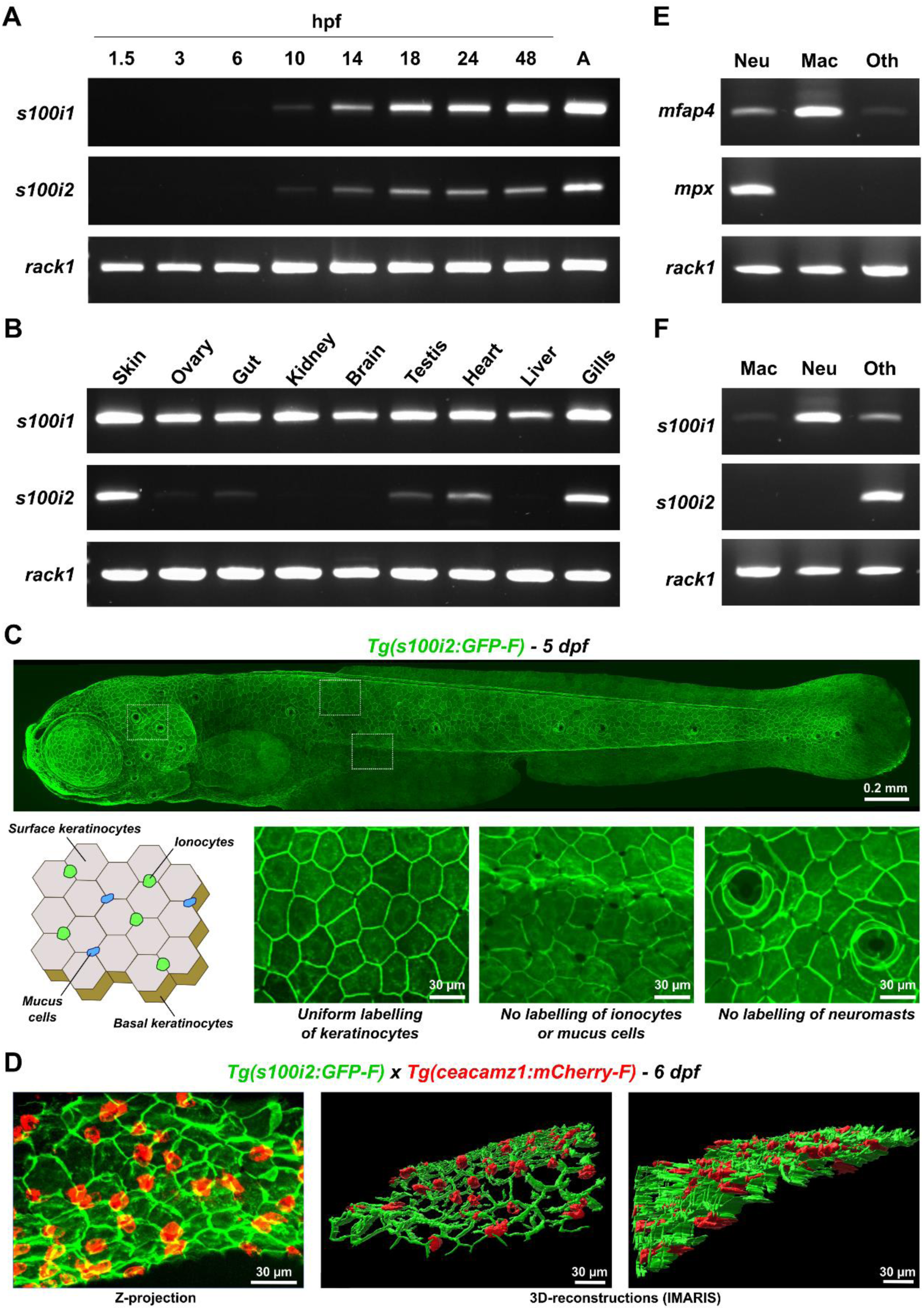
Spatio-temporal distribution of s100i1/i2 in zebrafish adults and larvae. **A.** RT-PCR analysis of s100i1/i2 expression during development (developmental stages indicated in hpf). “A” refers to adult stage (3 months old). Both genes are readily detected at 10 hpf. **B.** RT-PCR analysis of s100i1/i2 expression in various tissues of adult zebrafish, highlighting the distinct expression pattern of the two duplicate genes. **C.** Live imaging of a 5 dpf Tg(s100i2:GFP-F) zebrafish larva using spinning disk confocal microscopy. The upper panel shows a mosaic reconstruction of the whole larva using maximum projections from confocal z-stacks, highlighting the uniform GFP labelling on the whole epidermis. Lower panels display magnifications on different regions of the epidermis, revealing uniform labelling of all keratinocytes whereas intercalating cells like mucus cells, ionocytes or neuromasts remain unlabelled. A schematic representation of the cell composition of zebrafish skin epidermis at early larval stages is included to ease the identification of the different cell types. **D.** Z-projection of a portion of the ventral skin of a 6 dpf double-crossed Tg(s100i2:GFP-F) x Tg(ceacamz1:mCherry-F) zebrafish larva (left) and corresponding 3D-reconstruction using IMARIS (right). The S100i2-expressing keratinocytes (green) are located at the same level as Ceacamz1-expressing ionocytes (red), thus corresponding to surface keratinocytes. **E.** RT-PCR analyses using mfap4 and mpx specific primers to evaluate the efficiency of the FACS procedure in obtaining macrophage and neutrophil-specific RNA samples. **F.** RT-PCR analysis of s100i1/i2 expression in zebrafish myeloid cells. The housekeeping rack1 gene was used as control in all RT-PCR experiments.

Noteworthily, both genes are strongly expressed in the skin, a feature also shared by mammalian *s100* genes. To further specify where in the skin *s100i1*/*i2* are expressed, we set out to generate reporter lines, which only succeeded for *s100i2*. The resulting transgenic line, Tg(*s100i2:GFP-F*), drives the expression of a farnesylated version of the GFP protein under the control of a 1.5 kb fragment of the *s100i2* promoter region. As shown in Figure 8C, the GFP signal uniformly labels all keratinocytes from the zebrafish larval epidermis, whereas interstitial cells like ionocytes, mucus cells or neuromasts remain unmarked (Fig. 8C, zoomed images). Although simpler than the human skin, zebrafish epidermis contains two layers of keratinocytes, both at larval and adult stages: the superficial one, in direct contact with the external environment, and the basal one, lining up against the basement membrane and connecting the underlying dermis (Fig. 8C). To identify which layer(s) of keratinocytes strongly express *s100i2*, we crossed our reporter line with the Tg(*ceacamz1:mCherry-F*) line, that labels superficial HR-rich ionocytes [20]. As shown in Figure 8D, there is no overlapping of the green and red fluorescence signals in the larval progeny from this double cross, which confirms that ionocytes are not expressing *s100i2*. 3D-reconstructions using IMARIS software further show that the GFP-labeled keratinocytes are located at the same level as HR-rich ionocytes on zebrafish epidermis, i.e. they correspond to the upper layer of keratinocytes (Fig. 8D). Thus, basal keratinocytes do not seem to produce detectable levels of *s100i2*-driven GFP, at larval stages at least, suggesting that *s100i2* expression is restricted to surface keratinocytes. Finally, a hallmark of mammalian S100 proteins, especially the ones acting as pro-inflammatory alarmins, is their abundant expression in myeloid cells. To evaluate the presence of *s100i1*/*i2* transcripts in zebrafish macrophages and neutrophils, the two myeloid cell types most represented at larval stages, we used a protocol established in our laboratory that allows to obtain high quality RNA samples following FACS-assisted sorting of macrophage and neutrophil populations from dissociated zebrafish larvae [21]. For this purpose, we used double-fluorescent 5 dpf larvae from the crossing between reporter line Tg(*mfap4.1*:*mCherry-F*) (specifically labelling macrophages with red fluorescence) and reporter line Tg(*mpx*:*GFP*) (specifically labelling neutrophils with green fluorescence). Efficacy of the cell sorting was first validated by RT-PCR, using primers that amplify markers specific for macrophages (*mfap4* gene) or neutrophils (*mpx* gene). As seen in Figure 8E, *mfap4* transcripts are highly enriched in the RNA samples extracted from macrophages whereas *mpx* transcripts are only detected in the RNA samples extracted from neutrophils, thus validating our procedure. Evaluation of *s100i1*/*i2* expression in these cell types showed that *s100i1* is abundantly expressed in neutrophils and more faintly detected in macrophages, while *s100i2* is barely detected in any of these myeloid cells - observations in agreement with the single cell RNA seq data on expression of these two genes in zebrafish myeloid cells provided in the BASICz database (https://www.sanger.ac.uk/tool/basicz/; Supplementary Fig. S2) [22].

Altogether, these data reveal that S100i1/i2 also share features with mammalian S100s at the gene expression level, being abundantly expressed in tissues exposed to the environment, especially the skin, and in cell types that are also important reservoir for S100 proteins in mammals, such as keratinocytes and myeloid cells.

### Regulation of ictacalcin gene expression in disease context

In mammals, S100s contribute to many pathologies with deleterious outcome, as they often undergo up-regulation at the transcriptional level, which allows to sustain their pro-inflammatory action, thereby prompting unresolved inflammation and aggravated tissue damage. To assess whether such transcriptional control also occurs for zebrafish ictacalcins, we investigated their expression levels in representative zebrafish disease models of varying severity. First, we employed the tail fin amputation model on 3 dpf larvae, which generates a rapid but rather mild and short-lived inflammatory insult, as resolution of inflammation initiates within the first day following injury and full regeneration of the amputated fin fold occurs within 3-5 days [23]. In this model, we observe strong transcriptional up-regulation of pro-inflammatory cytokine genes such as *tnfa.a* or *il1b* within the first 2-3 hours following amputation (hpa), while the levels of mRNA transcripts for these genes rapidly drop down after 5 hpa, indicating the start of the resolution phase (Fig. 9A, upper graphs). In contrast, no significant modulation of *s100i1*/*i2* mRNA levels can be observed in this time window (Fig. 9A, lower graphs).

**Figure 9.**
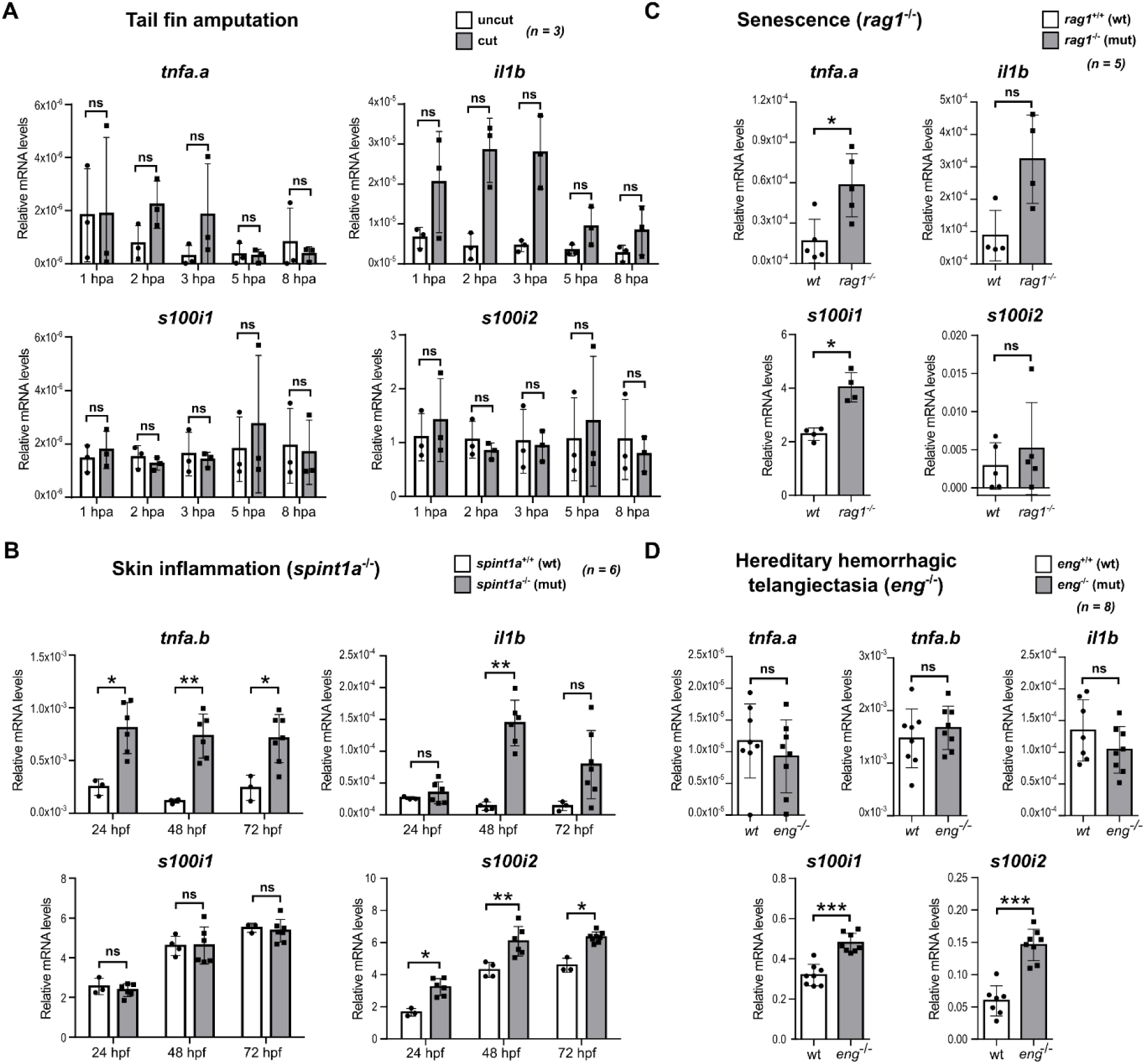
Transcriptional modulation of s100i1/i2 in various zebrafish models of inflammation and/or chronic disease. **A-D.** RT-qPCR analyses of s100i1/i2 gene expression as compared to housekeeping gene in the tail fin amputation model in 3 dpf larvae (**A**), in the spint1a^-/-^ model of skin inflammation in 1 to 3 dpf larvae (**B**), in the rag1^-/-^ model of senescence in 4 month old adults (**C**) and in the eng^-/-^ model of HHT in 1 month old juveniles (**D**). The relative mRNA levels of the classical pro-inflammatory markers tnfa.a, tnfa.b and il1b were also evaluated for comparison. The housekeeping rack1 and ef1a genes were used as control in all qPCR experiments. The number of independent replicates (n) is indicated on each panel (corresponding to number of independent batches of larvae for larval models or number of individuals for adult models). The statistical significance of the observed differences was analyzed using non-parametric Mann-Whitney test. P-values are reported as follow: *: 0.05 > p > 0.01; **: 0.01 > p > 0.001; ***: 0.001 > p > 0.0001; ****: p < 0.0001; ns: not significant.

We next moved to a more intense inflammatory context and investigated the *spint1a*^-/-^ mutant model, which results in strong skin inflammation with abundant neutrophilic infiltration and psoriasis-like phenotype with hyperproliferative keratinocytes [24,25]. In this model, the inflammatory burden is most intense within the first 3 days of life, healthy phenotype being restored at 5 dpf through gene compensation. Accordingly, we observe strong up-regulation of *tnfa.b* and *il1b* mRNA transcripts in mutant larvae at 48 and 72 hpf (as well as 24 hpf for *tnfa.b*) (Fig. 9B, upper graphs). Interestingly, while no modulation of *s100i1* gene expression can be detected, the levels of *s100i2* transcripts are significantly up-regulated at all time points investigated in the mutant larvae (Fig. 9B, lower graphs), revealing for the first time transcriptional control of a zebrafish *s100* gene in a larval model of sterile inflammation.

To evaluate whether such modulation may also occur in adult models of sterile inflammation, we moved to the *rag1^-/-^*mutant model of senescence, in which mutant fish lack adaptive immunity and therefore develop early signs of aging, as well as higher innate immune activity [26]. Such model is highly relevant for S100 biology as mammalian S100s are heavily involved in pathological inflammation in the aged brain [27]. Enhancement of immune activity at the transcriptional level was investigated in the primary lympho-myeloid organ of the fish, the head kidney, in 4 months old individuals - stage at which the worst inflammatory phenotype is noted. In accordance, both *tnfa.a* and *il1b* transcripts are enriched in the mutant fish at this stage, as compared to wild-type animals (Fig. 9C, upper graphs). *s100i1* also displays significant transcriptional up-regulation while *s100i2* does not show any statistically significant modulation (Fig. 9C, lower graphs).

Finally, we explored *s100i1*/*i2* modulation in the *eng^-/-^* mutant model recently developed in our laboratory that recapitulates key symptoms of hereditary hemorrhagic telangiectasia (HHT), a genetic condition causing arteriovenal malformations in humans [28]. It was previously demonstrated that the cardiovascular defects observed in zebrafish *eng^-/-^* mutants are associated with chronic hypoxia. On the other hand, the inflammatory status in *eng^-/-^* fish has not yet been assessed. Therefore, we first evaluated mRNA transcript levels for the control markers *tnfa* (both *tnfa.a* and *tnfa.b* isoforms) and *il1b* in 1 month old whole zebrafish. As shown in Figure 9D (upper graphs), none of these pro-inflammatory genes is significantly modulated in *eng^-/-^*individuals as compared to WT fish. In contrast, both *s100i1* and *s100i2* are significantly up-regulated in *eng^-/-^* mutants (Fig. 9D, lower graphs).

Altogether, these data demonstrate that teleost-specific *s100* genes undergo transcriptional up-regulation in disease contexts associated with sustained inflammation or chronic hypoxia, likewise mammalian S100s.

## Discussion

We here report the very first and detailed structural characterization of two teleost-specific S100 proteins, ictacalcins S100i1 and S100i2 from *Danio rerio*. High resolution crystallographic structures of the proteins in various ionic states allowed us to demonstrate that although not orthologous to mammalian S100s, they share all their structural features, including the two non-equivalent EF-loop motifs per protomer and the quaternary organization into homodimers. We also confirmed that they bind two calcium ions per protomer with micromolar affinities, in agreement with the values reported for mammalian S100s [18]. Furthermore, the conformational changes we observed upon occupation of the EF-loops of S100i1/i2 by sodium or magnesium led us to propose that they undergo calcium-dependent conformational activation, likewise mammalian S100s. This model is corroborated by a recent study from the Harms group where they evidenced calcium-induced changes in the secondary and tertiary structure of zebrafish S100i1 upon Ca^2+^ addition, using far-UV circular dichroism and intrinsic fluorescence [11].

We also showed that neither ictacalcins bind to zinc, as a surrogate for transition metals. In mammals, several S100 proteins can chelate metal ions, these are mainly the ones involved in nutritional immunity like calprotectin, S100A7 or S100A12. Absence of metal chelation by S100i1/i2 therefore suggests that they do not possess direct antimicrobial activity. This was recently confirmed for S100i1, which failed to restrict bacterial growth of *Staphylococcus epidermidis* and zebrafish-derived *Aeromonas* or *Vibrio* strains in *in vitro* assays, in contrast to human calprotectin [11]. These findings are somewhat surprising since many of the zebrafish *s100* genes, including *s100i1* and *s100i2*, were found strongly up-regulated following infection by viral or bacterial fish pathogens [29–31]. Furthermore, in other teleost species, ictacalcin genes were also found significantly up-regulated both at the gene and protein levels on mucosal surfaces. For instance, upon infection of salmon (*Salmo salar*) with the ectoparasite *Lepeophtheirus salmonis*, *s100i1* appeared significantly up-regulated in the skin mucus, suggesting a role in mucosal immunity against this pathogen [32]. These are therefore good indicators that if not through direct bactericidal action, ictacalcins may also contribute to antimicrobial defenses through indirect mechanisms, for instance by modulating immune and inflammatory responses against infectious agents.

In line with this idea, our results highlight the strong expression of ictacalcins in tissues and cells with immune functionalities. At adulthood, *s100i1* and *s100i2* are both found highly expressed in the skin and gills, and *s100i1* is also present in the intestine - three major organs directly in contact with the microbial-rich aquatic milieu and providing important mucosal defenses towards external aggressions. Furthermore, *s100i1* is abundantly detected in neutrophils, the first immune responders arriving at sites of injury. This strongly parallels the behavior of mammalian S100s, as the most active contributors to innate immunity in humans, S100A7 and the calgranulins S100A8, S100A9, and S100A12, share similar tissue and cell distribution, being predominantly expressed in the respiratory and digestive tracts (A7/A8/A9), in the skin (A7/A8/A9) and in the bone marrow and lymphoid tissues (all four) [33]. Furthermore, the mammalian S100A8/A9 heterodimer constitutes up to 45% of the pool of cytosolic proteins in neutrophils [34]. Noteworthily, we detected *s100i1*/*i2* expression early during embryogenesis (10 hpf) and transcriptomic studies by Zhang *et al* [29] showed that *s100i1*/*i2* are amongst the zebrafish *s100* genes most highly expressed during the first days of life, with mRNA levels being two to five times higher than those of housekeeping gene *ef1a*. Such early and abundant presence again suggests that S100i1/i2 may be required for protective function in the developing embryo, even before the appearance of functional innate immune cells.

Interestingly, human skin epidermis constitutes an important reservoir for S100 proteins, as sixteen distinct S100s are found in this tissue [35]. Their genes are all clustered together in what is known as the Epidermal Differentiation Complex (EDC), which also encodes other gene families essential for normal epidermal differentiation [35,36]. Epidermal S100s exert distinct functions depending on the layer they occupy. Basal ones play important roles in epidermal development and cell differentiation, promoting keratinocyte proliferation and assisting the renewal of epidermal cells. S100s from the middle layer have a predominant role in controlling the trafficking of membrane receptors, allowing communication between the different layers and thereby enabling epidermal growth. S100s from the upper layers (granular and upper spinous layer) contribute to keratinocyte differentiation in healthy skin and regulate calcium homeostasis [35]. We here demonstrated that zebrafish skin also expresses at least two S100 members. Although their synteny is not conserved with that of mammalian *s100* genes, *s100i1* and *s100i2* cluster with other *s100* genes (*s100a10b*, *s100w*) on zebrafish chromosome 16, as well as with keratinocyte-associated genes (*krtcap2*) or genes involved in signaling during epidermal differentiation (*efna1b*, *efna3b*), which suggests that S100i1/i2 may also contribute to keratinocyte biology and/or epidermal development. In mammals, S100s from the upper epidermal layer also heavily contribute to the first-line defense against pathogens and to the maintenance of epithelial integrity, either through direct antimicrobial activity or by activation of keratinocytes and infiltrating immune cells that in turn produce cytokines and chemokines to counteract the damage. The predominant expression of S100i2 we observe in all epidermal keratinocytes from the upper layer, similar to that already reported for S100i1 [13], therefore suggests possible conserved functions in mucosal immunity, especially since surface keratinocytes from larval epidermis also contain high amounts of pro-inflammatory cytokines like Il-1b [20,37].

With respect to their alarmin function, another hallmark of mammalian S100s is their transcriptional up-regulation following prolonged injury, meant to sustain continuous feeding of the pro-inflammatory response. While up-regulation of zebrafish *s100* genes was previously demonstrated in several adult models of infection, here we provide the first examples of transcriptional control of teleost-specific *s100* gene expression in sterile disease contexts, both in adults and larvae. Importantly, we show that injury models associated with short-lived and mildly intense inflammatory burden, like the tail fin amputation model, do not elicit transcriptional modulation of *s100i1/i2* genes. In contrast, more sustained inflammatory damage, like in the *spint1a^-/-^* model of skin inflammation or in the *rag1^-/-^* model of senescence, promote their differential expression. Interestingly, the two ictacalcin genes display distinct modulation in these models. Keratinocyte-abundant *s100i2* is up-regulated in the *spint1a^-/-^* model, linked to abnormal keratinocyte behavior, whereas broadly expressed *s100i1* is up-regulated in the head kidney of *rag1^-/-^* mutants. Differential tissue distribution and transcriptional modulation of *s100i1*/*i2* thus provides another example of two duplicate zebrafish genes that evolved independently following the third whole genome duplication event [38], in order to acquire specific expression features and so that their gene products may exert distinct biological roles rather than functioning as redundant proteins.

Our evidence that *s100i1*/*i2* get transcriptionally modulated during long-lasting inflammatory insult again establishes a strong parallel with mammalian S100s and hint for a possible role of zebrafish ictacalcins in the development of deleterious inflammatory phenotypes. This may be particularly true for skin pathologies in which mammalian S100s are strongly implicated, like skin cancers, dermatitis or psoriasis [39]. Psoriasis for example is characterized by an hyperproliferation of keratinocytes and an abnormal infiltration of T-cells, dendritic cells, macrophages, and neutrophils in the skin [40]. Keratinocyte-derived S100A7 and S100A15 are strongly up-regulated in human psoriatic skin and drive the chemotactic recruitment of immune cells together with the sustained production of pro-inflammatory cytokines, further potentiating the inflammatory cascade [41]. Zebrafish *spint1a^-/-^* mutant provides a highly relevant model of this pathology, with hyperproliferative behavior of keratinocytes, aberrant immune infiltration and epidermal aggregates mimicking psoriatic lesions [42]. The up-regulation of *s100i2* we observe in this model, concomitant to that of pro-inflammatory markers *tnfa.b* and *il1b*, thus strengthens the parallel with the human pathology and calls for a deeper evaluation of the role possibly played by zebrafish ictacalcins in the propagation of the psoriatic inflammatory symptoms.

Finally, we also evidenced transcriptional up-regulation of *s100i1*/*i2* in the *eng^-/-^* model, which may not be characterized by an underlying inflammatory context, since we did not detect significant up-regulation of classical pro-inflammatory markers. This model is associated with chronic hypoxia [28], a condition often encountered in a cancer context in humans and to which S100 proteins have already been shown to respond and/or contribute [43]. Indeed, hypoxic conditions can increase the production of several S100s, like S100A4 or the highly pro-inflammatory S100A8/A9 [44,45]. Conversely, S100s and their downstream signaling can directly control hypoxia-dependent processes, notably through the modulation of HIF-1α expression [46]. Many of these processes are dependent on the receptor for advanced glycation end-products (RAGE), the main receptor for S100s in mammals, which is absent in zebrafish. A direct implication of S100i1/i2 in modulating hypoxic conditions in the *eng^-/-^* model - and more generally in disease contexts in zebrafish - thus remains an open question, but other S100 receptors highly conserved in zebrafish, such as Toll-like receptors or yet to discover teleost-specific S100 receptors, may relay such function.

In conclusion, our study demonstrates that although teleost-specific S100i1/i2 do not share strict orthology with any mammalian S100, they exhibit highly conserved structural and biochemical features at the protein level, similar tissue distribution and parallel gene modulation during inflammation or hypoxia. They may thus also play a role in regulating immune and inflammatory responses in chronic disease contexts. Defining the exact nature and extent of this role evidently deserves further investigation, including the generation of KO lines or loss-of-function mutants to be used in the zebrafish inflammatory models we already explored.

## Methods

### Expression and purification of recombinant S100i1 and S100i2 from *Danio rerio*

The full-length ORFs of zebrafish S100i1 and S100i2 proteins were PCR-amplified from *Danio rerio* genomic DNA and cloned in frame after the sequence coding for Tobacco Etch Virus (TEV) protease cleavage site in pETM11 vector (EMBL vector collection), using restriction-free cloning [47]. CterW mutants of S100i1 and S100i2 were generated from these constructs using PCR-based site directed mutagenesis with High Fidelity Phusion DNA Polymerase (New England Biolabs) and anti-complementary oligonucleotides bearing the Trp insertion. For expression of S100i1 and S100i2, the pETM11 constructs were transformed into heat-competent *E. coli* BL21 (DE3). Cells were grown at 37°C in 2xYT medium supplemented with 50 μg/ml of kanamycin, until they reached an exponential phase of growth. Protein expression was induced by adding 1 mM IPTG and further incubating overnight at 18°C. The recombinant proteins were then purified to homogeneity from *E. coli* soluble cell extracts, using our established 3-step procedure [48] consisting of a first Nickel-affinity chromatography (Ni-NTA), followed by TEV cleavage and counter-purification on Ni-NTA, and a final step of size exclusion chromatography (SEC) performed on a 24 ml Superdex 75 Increase 10/300 GL column (Cytiva) equilibrated with 20 mM Tris-HCl pH 7.5, 100 mM NaCl.

### Crystallization and data collection

Purified S100i1/i2 proteins in buffer 20 mM Tris-HCl pH 7.5, 100 mM NaCl were concentrated to 10 mg/ml and set to crystallize either in apo form or after supplementation with 5 mM CaCl_2_ (5 eq), using the sitting-drop vapor diffusion method in 48- or 96-well plates (Swissci) and commercial crystallization screens (Molecular Dimensions Ltd). Crystal forms 1 and 2 of S100i1 appeared at 291K over a reservoir containing 0.2M Li sulfate, 0.1M Tris-HCl pH 8.5, 25% PEG 4000 or 0.2M Na acetate, 0.1M Tris-HCl pH 9.5, 35% PEG 4000, respectively, while S100i2 crystals grew at 291K using 0.2M MgCl_2_, 0.1M Tris HCl pH 8.5, 30% PEG 4000 as crystallization solution.

Prior to data collection, S100i1 crystals were soaked into a cryoprotective solution corresponding to the crystallization condition supplemented with 10% (crystal form 1) or 20% (crystal form 2) glycerol before flash-freezing into liquid nitrogen. S100i2 were directly flash-frozen into liquid nitrogen. Complete X-ray diffraction datasets were collected at a wavelength of 1.0 Å on the X06DA beamline at the Swiss Light Source facility (SLS, Switzerland).

### Structure determination and refinement

Datasets were processed with XDS [49]. Both crystal forms of S100i1 belonged to space group P2_1_ while S100i2 crystallized in P2 space group. The structures of Na^+^-S100i1 and Mg^2+^-S100i2 were determined by molecular replacement (MR) using automated model search with Balbes [50]. Interestingly, although S100i1 and S100i2 share 87% sequence identity, the structural models from two distinct S100 proteins were used for MR search, namely that of human Ca^2+^-bound S100A2 [51] for S100i1 and that of human Ca^2+^-bound S100A4 [52] for S100i2. The resulting S100i1 model was then employed to phase the X-ray data from apo S100i1, using MR in Phaser [53]. All three models were refined by alternating cycles of manual rebuilding in Coot [54] and cycles of energy minimization in PHENIX.REFINE [55], including refinement of individual isotropic Atomic Displacement Parameters (ADP) and Translation– Libration–Screw (TLS) parameterization (Table 1). S100i1 crystal form 2 and S100i2 crystals showed strong translational non-crystallographic symmetry (tNCS), which yielded final R and R_free_ values much higher than expected for the resolution. Nevertheless, the electron density maps were overall well defined and allowed to accurately rebuild all three structural models, except in a few loop regions showing more flexibility. The quality of the final models was assessed with Molprobity [56]. Atomic coordinates and structure factors have been deposited in the Protein Data Bank under accession codes 9IN2 (apo S100i1), 9HYG (Na^+^-bound S100i1) and 9I1G (Mg^2+^-bound S100i2).

### Sequence and structural analyses

Sequence alignments were done in CLUSTAL OMEGA [57] and coloring according to conservation was done in ALINE [58]. Analysis of S100 dimeric assemblies and interfaces was performed in PISA [14]. 3D-modeling of Ca^2+^-bound S100i1 and S100i2 was performed with AlphaFold3 server [59]. All figures were made with the Pymol Molecular Graphics System (version 215 0.99rc6, DeLano Scientific LLC).

### Isothermal titration calorimetry (ITC)

For ITC experiments, S100i1 and S100i2 proteins were purified as described previously except that the last step on SEC was performed either in Buffer 1 (20 mM Tris pH 7.5, 100 mM NaCl; for calcium titrations) or in Buffer 2 (20 mM Tris pH 7.5, 100 mM NaCl, 2 mM CaCl_2_; for zinc titrations). Frozen proteins were thawed on ice before use. ITC experiments were performed in a MicroCal PEAQ-ITC (Malvern Panalytical Ltd, UK) at 25 °C. The first injections were 0.4 μL, followed by 19 x 2 μL injections with a stirring rate of 750 rpm. To achieve Ca^2+^ binding saturation, a second series of injections were performed and the resulting ITC data files were concatenated into a single data file for analysis. S100i1 and S100i2 were used at 40 μM and 42-60 µM in the ITC cell, respectively; 0.4-0.8 mM CaCl_2_ or 0.6-2.0 mM Zn acetate solutions were used in the syringe. ITC data were integrated and baseline corrected using NITPIC [60]. The integrated data were globally analyzed in SEDPHAT [61] using a model with two symmetric binding sites (macroscopic K_D_) or two non-symmetric sites (microscopic K_D_). We also included a floating “fraction incompetent” parameter to capture uncertainty in the relative protein and Ca^2+^ concentrations. Thermogram and binding figures were plotted in GUSSI [62].

### Zebrafish lines, maintenance and ethics

All zebrafish experiments described in the present study were conducted by authorized staff in compliance with the European Union guidelines for handling of laboratory animals (2010/63/EU Directive) and the ARRIVE guidelines [63]. Breeding and maintenance of adult fish were performed at the ZEFIX-LPHI (CNRS, University of Montpellier, Montpellier, France; license number CEEA-LR-B34–172–37) and University of Murcia (CARM approval number #A13220914) fish facilities, according to the local animal welfare standards approved by the Direction Sanitaire et Vétérinaire de l’Hérault, the Comité d’Ethique pour l’Expérimentation Animale of Languedoc-Roussillon, the French Ministère de l’Enseignement Supérieur de la Recherche et de l’Innovation under reference APAFIS #36309-2022040114222432 V2 and by Comunidad Autónoma de la Región de Murcia, Dirección General de Ganadería, Pesca y Acuicultura.

Fish and embryo maintenance, staging and husbandry were as described previously [20]. Experiments were performed using AB, *golden* [64] or *casper* [65] zebrafish strains (ZIRC) along with the following transgenic lines: Tg(*mfap4.1:mCherry-F*) ump6Tg to label macrophages [66], Tg(*mpx:GFP*) i114Tg to label neutrophils [67], Tg(*ceacamz1:mCherry-F*) ump9Tg to label epidermal HR-ionocytes [20], Tg(*s100i2:GFP-F*) ump16Tg (this study) to visualize the transcriptional expression of *s100i2*, and the following mutant lines corresponding to loss-of-function for the targeted genes: *spint1a*^hi2217Tg/hi2217Tg^ or *spint1a^-/-^*(Serine Peptidase Inhibitor, Kunitz Type 1a gene) [24], *eng^-/-^* (Endoglin gene) [28] and *rag1^t26683/t26683^*or *rag1^-/-^* (Recombination activating gene 1) [26,68]. Zebrafish embryos were obtained from pairs of adult fish by natural spawning and raised at 28◦C under 14:10 hours light/dark cycle. All experimentations on live larvae were performed under anesthesia with tricaine (ethyl 3-aminobenzoate). When required, euthanasia was achieved using an overdose of tricaine (500 mg/L).

### FACS-sorting of macrophages and neutrophils

Macrophage and neutrophil cell populations were sorted from dissociated zebrafish larvae using FACS as previously documented [21]. Briefly, for one batch, 120 larvae obtained from the crossing between Tg(*mfap4.1:mCherry-F*) and Tg(*mpx:GFP*) lines, and selected based on double-positive fluorescence, were dissociated at 5 or 6 dpf using our established enzyme-free cell dissociation procedure. After final resuspension in buffer 0.9X Dulbecco’s Phosphate-Buffered Saline (DPBS) supplemented with 2% heat-inactivated Fetal Bovine Serum (FBSi) and 2 mM EDTA, SYTOX™ Red Dead Cell Stain was added in the sample (5 nM final concentration) to label non-viable cells. Live mCherry-positive, GFP-positive and double negative cell populations were then sorted by FACS on a BD FACSAria™ IIu cell sorter on the Montpellier RIO Imaging microscopy platform (MRI, Biocampus, Montpellier), using gating strategy and fluorescence detection parameters as previously described [21]. Sorted cells were directly collected into RNase/DNase-free 1.5 ml microfuge tubes containing 100 µl of lysis buffer from the QIAGEN RNeasy Micro kit (RLT buffer supplemented with 1% v/v β-mercaptoethanol). The samples were vigorously vortexed and then immediately flash-frozen in liquid nitrogen for storage at -80°C until proceeding with RNA extraction.

### Total RNA extraction

For *s100i1*/*i2* expression studies at larval stages, 25-50 embryos/larvae from the desired zebrafish lines were pooled and their total RNA was isolated with the NucleoSpin RNA Plus kit (Macherey-Nagel). Expression studies during development were performed on dechorionated WT AB larvae collected at various developmental stages (1.5 to 48 hpf). Total RNA for adult control was extracted from 2-3 adult fish (3 months old) using NucleoZOL protocol (Macherey-Nagel). Expression studies in the tail fin amputation and *spint1a^-/-^* models were conducted on tail-fin amputated versus intact 3 dpf WT AB larvae or on *spint1a^-/-^* versus *casper* (*spint1a^+/+^*) 24/48/72 hpf larvae, respectively. RNA samples for expression studies in adult tissues/organs were obtained previously [20]. RNA samples for expression studies in *rag1^-/-^* (head kidney of 4 months old zebrafish), *eng^-/-^*(1 month old whole fish) and wild-type siblings (*rag1^+/+^*, *casper* background; *eng^+/+^*, AB background) were from previous reports [26,28]. Macrophage-specific, neutrophil-specific and double-negative total RNAs were extracted from FACS-sorted samples using the RNeasy Micro kit (QIAGEN) and RNA sample quality and concentration was assessed using the RNA 6000 Pico kit (Agilent) on an Agilent 2100 Bioanalyzer System (qPhD platform, Montpellier). All expression studies were performed on at least three independent replicates, for each time point/stage/condition investigated.

### Semi-quantitative and quantitative PCR

For primer specificity check, the full-length coding sequences of all fourteen zebrafish *s100* genes were amplified from *D. rerio* cDNA and cloned into pETM11 vector using RF-cloning. For expression studies, equal amounts of total RNAs (500 ng) were reverse-transcribed using Oligo(dT) primers and M-MLV reverse transcriptase (ThermoFisher). For RNA samples extracted from FACS-sorted cells, only 4 ng of total RNAs were used for reverse transcription due to low yields of RNA. Semi-quantitative PCR analyses (RT-PCRs) were performed with an amount of cDNA equivalent to 5 ng total RNA or 0.1 ng total RNA for expression studies in myeloid cells or on 10 ng of purified pETM11-S100 plasmid, using Phusion DNA polymerase (New England Biolabs) and specific sets of primers for each gene as listed in Table S1.

Quantitative PCR (qPCR) reactions were carried out using an amount of cDNA equivalent to 0.5 ng total RNA in SYBR^®^ Green mix (SensiFast™ SYBR^®^, Meridian BioScience) in the presence of 600 nM of each primer (as indicated in Table S1). Reaction mixes were assembled in 384-well plates with help of a Labcyte Echo 525 Liquid Handler and qPCRs were ran in triplicates using a LightCycler® 480 II thermal cycler (Roche) available at the Montpellier GenomiX’s High-throughput qPCR facility (MGX, Biocampus, Montpellier). Data were analyzed with the LC480 software and results are presented as relative target gene mRNA expression as compared to reference gene (*rack1* or *ef1a*).

### Generation of the Tg(*s100i2:GFP-F*) fluorescent reporter line

The Tg(*s100i2:GFP-F*) transgenic reporter line was generated following previously reported procedures [20,37]. A 1.5 kb fragment of the *icn2* (*s100i2*) promoter was amplified from zebrafish genomic DNA using 5’-CTTCAAAATGACAATTCCCATACTTG-3’ and 5’-CATGTTTCTGGCTGTAAAAAAAAGAC-3’) as forward and reverse primers. This fragment, which terminates at the *icn2* AUG start codon, was inserted just upstream of the reading frame of farnesylated GFP protein (GFP-F) in the I2BN-modeGFP transgenesis vector, using restriction-free cloning. This vector, derived from pBluescript, harbors two I-SceI homing sites [69]. The resulting plasmid was injected into one-cell stage embryos (wild-type AB zebrafish line) together with the I-SceI meganuclease. F0-microinjected embryos were screened for transgene expression at 1 and 2 dpf, and stable lines were established. The line has been registered as ump16Tg in ZFIN.

### Confocal microscopy and image analysis

Imaging of 5 and 6 dpf larvae from the Tg(*s100i2:GFP-F*) reporter line or from double-cross with the Tg(*ceacamz1:mCherry-F*) line was done by confocal microscopy. Prior to live imaging, larvae were anesthetised with 0.016% (w/v) tricaine and immobilized in 1% low-melting point agarose in 35 mm glass-bottom FluoroDish plates (World Precision Instruments, UK). Fluorescence image stacks were acquired at 28°C using a spinning disk Nikon Ti Andor CSU-W1 confocal microscope mounted with 20x/0.75 or 40x/1.15 Water objectives and an ANDOR Neo sCMOS camera (lasers 488 nm for GFP and 561 nm for mCherry). Images were processed with Fiji (Image J software) and compressed into maximum intensity Z-projections. 3D-reconstructions were generated with IMARIS software (Oxford Instruments, https://imaris.oxinst.com). Image acquisition and processing were performed on the Montpellier RIO Imaging microscopy platform (MRI, Biocampus, Montpellier).

### Statistical analyses

Expression studies were designed to generate experimental groups of comparable size, defined from preliminary experiments. No inclusion/exclusion criteria of data were applied. Relative mRNA expression levels were represented as mean ± standard deviation. GraphPad Prism v8.3.0 (San Diego, CA, USA) software was used to construct graphs and analyze data. Non-parametric Mann-Whitney test was employed for statistical analyses and p-value p<0.05 was considered as threshold for the statistical significance of differences between groups. The number of independent experiments and the p-value definitions are indicated in the figure legends where appropriate.

## Author Contributions

LY designed the study; LY, MNC, VM, EL and SDT planned the experiments; LH, TP, MD, CBP, JFRV, SDT, CBi, CBu, CG, JG, EL and LY performed experiments; LY, MNC, VM, EL, and SDT provided student or technical staff supervision; LH, TP, MD, CBi, and LY analyzed the data and prepared the figures; LY wrote the original manuscript draft of the paper; all authors discussed the results and contributed to the final version of the manuscript.

## Acknowledgements

We thank the beamline staff at SLS for providing technical assistance during data collection. We are grateful to Victor Goulian from the ZEFIX-LPHI Aquatic model facility (University of Montpellier) and Pedro Martínez from Zebrafish Animal Facility (University of Murcia) for their technical support on zebrafish line maintenance. We also would like to thank Philippe Clair from the Montpellier GenomiX’s High-throughput qPCR facility (MGX, Biocampus), Myriam Boyer-Clavel and Stéphanie Viala from the MRI-IGMM cytometry facility (MRI, Biocampus), as well as Vicky Diakou and Elodie Jublanc from the MRI-DBS imaging facility (MRI, Biocampus) for their precious help and advice during data acquisition and analysis. We also acknowledge the imaging facility BioCampus Montpellier Ressources Imagerie (MRI), member of the national infrastructure France-BioImaging supported by the French National Research Agency (ANR-10-INSB-04, ’Investments for the future’). This work was supported by a grant from the European Community’s H2020 Program Marie-Curie Innovative Training Network INFLANET (https://inflanet.eu/): Grant Agreement nr. 955576. Access to the ITC platform at CBI-IGBMC was supported by the French Infrastructure for Integrated Structural Biology (FRISBI) ANR-10-INBS-0005.

