## Supplementary Information for "Teleost-specific ictacalcins exhibit similar structural organization, cation-dependent activation and transcriptional regulation as human S100 proteins"

### CONTENTS


| | |
|---|---|
| **Supplementary Figure S1** | SEC elution profiles of the CterW mutants of S100i1 and S100i2 |
| **Supplementary Figure S2** | Single blood cell gene expression profiles for *s100i1* and *s100i2* |
| **Supplementary Table S1** | List of primers used for semi-quantitative and qPCRs |

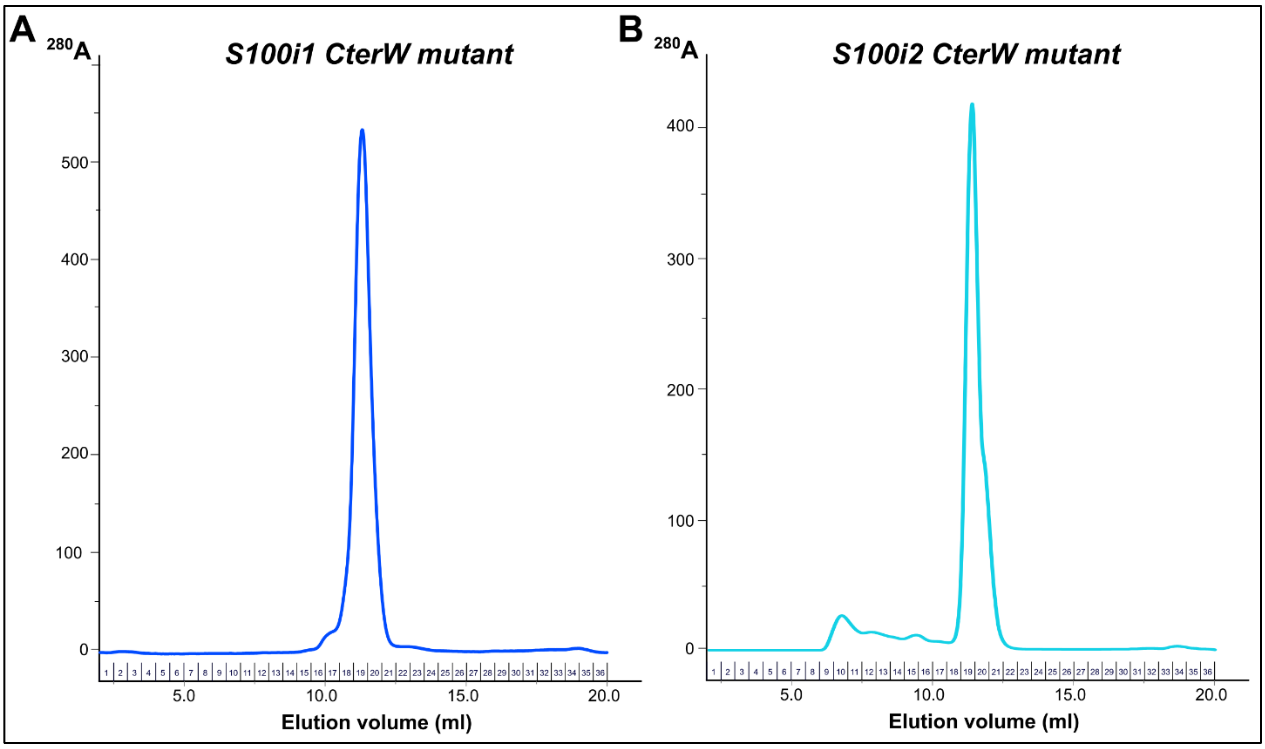


**Supplementary Figure S1. SEC elution profiles of the CterW mutants of S100i1 and S100i2.** Elutions were performed on a Superdex 75 Increase column (24 ml; Cytiva) equilibrated in 20 mM Tris-HCl pH 7.5, 100 mM NaCl. Both proteins eluted predominantly as homodimers, likewise WT proteins.

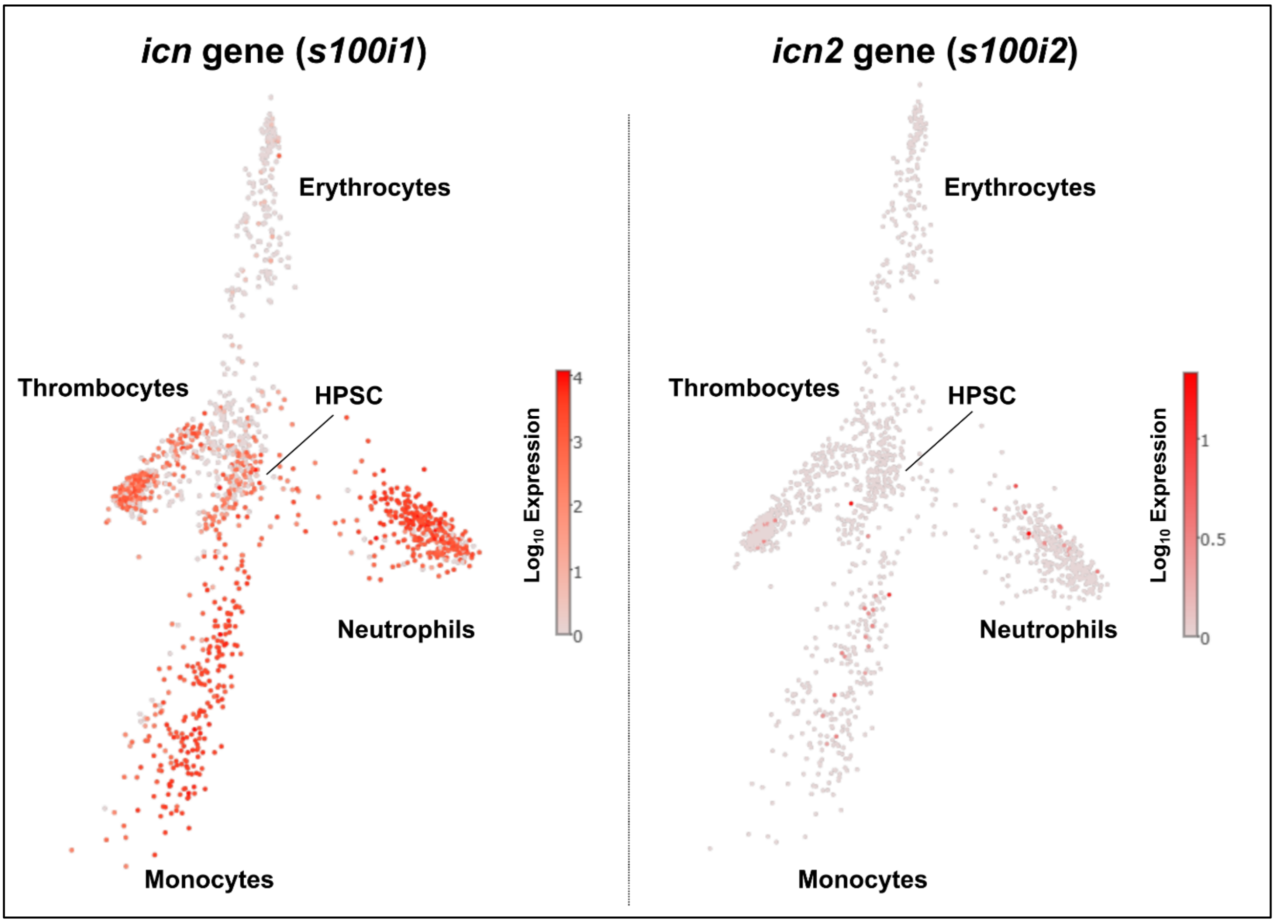


**Supplementary Figure S2. Single cell gene expression profiles for *s100i1* and *s100i2* in zebrafish blood cells according to BASiCz database.** Data are represented as diffusion maps covering the whole spectrum of blood cells in zebrafish (thrombocytes, erythrocytes, macrophages, neutrophils and hematopoietic stem cells or HPSC). Expression is represented as $Log_{10}$ fold and colored with gradient from white (no expression) to red (high expression).

| Gene | Primer sequence | Application |
|---|---|---|
| *s100i1* | Forward: 5’-GAACCACCATGGCTACGTCAG-3’<br>Reverse: 5’-CTGTAAGGACACTTATTTTTTTCCTGTG-3’ | *Semi-quantitative PCR* |
| | Forward: 5’-GTCCAAAGGCGAGCTGAAGGAAC-3’<br>Reverse: 5’-CTGTAAGGACACTTATTTTTTTCCTGTG-3’ | *qPCR* |
| *s100i2* | Forward: 5’-GCCAGAAACATGAGTGGCAAAATG-3’<br>Reverse: 5’-CCATAAGAACACACTTATTTTCCTTTTTGG-3’ | *Semi-quantitative PCR* |
| | Forward: 5’-GCCAGAAACATGAGTGGCAAAATG-3’<br>Reverse: 5’-CAGACACAACCATAAGAACACACTTATTTTCC-3’ | *qPCR* |
| *mfap4* | Forward: 5’-GCTGTTGAGGAGAGAGTGAGAAGATG-3’<br>Reverse: 5’-GTCAGCTGGTAGAGGTTCTCTAGTC-3’ | *Semi-quantitative PCR* |
| *mpx* | Forward: 5’-GGCTGCTGTTGTGCTCTTTCAATG-3’<br>Reverse: 5’-GGTTTGAGCTTCACAGCCTGTCATAC-3’ | *Semi-quantitative PCR* |
| *il1b* | Forward: 5’-TGGACTTCGCAGCACAAAATG-3’<br>Reverse: 5’-GTTCACTTCACGCTCTTGGATG-3’ | *qPCR* |
| *tnfa.a* | Forward: 5’-TTCACGCTCCATAAGACCCA-3’<br>Reverse: 5’-CCGTAGGATTCAGAAAAGCG-3’ | *qPCR* |
| *tnfa.b* | Forward: 5’-CGAAGAAGGTCAGAAACCCA-3’<br>Reverse: 5’-GTTGGAATGCCTGATCCACA-3’ | *qPCR* |
| *rack1* | Forward: 5’-CCTCGCCAAAATGACCGAGC-3’<br>Reverse: 5’-GGTGTACTTGCAGACTCCCAG-3’ | *Semi-quantitative PCR* |
| | Forward: 5’- CAGTTTGCTCTGTCTGGATCCTGG-3’<br>Reverse: 5’- GTGTACTTGCAGACTCCCAGAGTG-3’ | *qPCR* |
| *ef1a* | Forward: 5’-TTCTGTTACCTGGCAAAGGG-3’<br>Reverse: 5’-TTCAGTTTGTCCAACACCCA-3’ | *qPCR* |

**Supplementary Table S1. List of primers used for semi-quantitative and qPCRs.**
